# Engineering the biological conversion of formate into crotonate in *Cupriavidus necator*

**DOI:** 10.1101/2023.03.14.532570

**Authors:** Florent Collas, Beau B. Dronsella, Armin Kubis, Karin Schann, Sebastian Binder, Nils Arto, Nico J. Claassens, Frank Kensy, Enrico Orsi

## Abstract

To advance the sustainability of the biobased economy, our society needs to develop novel bioprocesses based on truly renewable resources. The C1-molecule formate is increasingly proposed as carbon and energy source for microbial fermentations, as it can be efficiently generated electrochemically from CO_2_ and renewable energy. Yet, its biotechnological conversion into value-added compounds has been limited to a handful of examples. In this work, we engineered the natural formatotrophic bacterium *C. necator* as cell factory to enable biological conversion of formate into crotonate, a platform short-chain unsaturated carboxylic acid of biotechnological relevance. First, we developed a small-scale (150-mL working volume) cultivation setup for growing *C. necator* in minimal medium using formate as only carbon and energy source. By using a fed-batch strategy with automatic feeding of formic acid, we could increase final biomass concentrations 15-fold compared to batch cultivations in flasks. Then, we engineered a heterologous crotonate pathway in the bacterium *via* a modular approach, where each pathway section was assessed using multiple candidates. The best performing modules included a malonyl-CoA bypass for increasing the thermodynamic drive towards the intermediate acetoacetyl-CoA and subsequent conversion to crotonyl-CoA through partial reverse β-oxidation. This pathway architecture was then tested for formate-based biosynthesis in our fed-batch setup, resulting in a two-fold higher titer, three-fold higher productivity, and five-fold higher yield compared to the strain not harboring the bypass. Eventually, we reached a maximum product titer of 148.0 ± 6.8 mg/L. Altogether, this work consists in a proof-of-principle integrating bioprocess and metabolic engineering approaches for the biological upgrading of formate into a value-added platform chemical.

## 1. Introduction

The current market volume of biobased chemical production is estimated in the order of ∼US$ 80 billion, with an expected annual growth rate of ∼10% (Nielsen et al., 2022). Despite this encouraging prediction, most economically feasible bioprocesses are limited to the production of fine chemicals (*i*.*e*., pharmaceuticals or food ingredients) from sugar feedstocks or agricultural waste (Jullesson et al., 2015; Nielsen et al., 2022; Paulino et al., 2021; Schempp et al., 2018). Nevertheless, these plant-based feedstocks are limited in availability and their use could compete with the food-supply chain.

As promising alternative, CO_2_ itself is a virtually free and unlimited feedstock which can be converted *via* renewable electricity to several C1 compounds like CO, formate, and methanol. These can serve both as carbon and energy sources in microbial cultivations (Satanowski and Bar-Even, 2020). In particular, methanol and formate hold potential as microbial feedstocks as they are soluble and can be easily handled and transported (Claassens et al., 2019; Cotton et al., 2020; Satanowski and Bar-Even, 2020). Although methanol has long been investigated for supporting microbial production (Dijkhuizen et al., 1985; Wang et al., 2020; Zhu et al., 2020), formate has emerged in the past decade as attractive feedstock for the bioeconomy (Bar-Even et al., 2013; Yishai et al., 2016). In fact, the latter can be generated through one-step electrocatalysis with high efficiencies and with long operational durations (Stöckl et al., 2022; Yang et al., 2020; Zheng et al., 2021). These advantages motivated the interest within the synthetic biology community in establishing synthetic formatotrophic growth, which has been implemented in the model organism *Escherichia coli via* different assimilation routes: the Calvin Benson Bassham (CBB) cycle (Gleizer et al., 2019), the reductive glycine pathway (rGlyP) (Bang et al., 2020; Kim et al., 2020), and the serine threonine cycle (Wenk et al., 2022). Owing to its linear architecture and low ATP cost, the rGlyP has also been functionally implemented in other non-traditional bacteria, such as *Cupriavidus necator* (Claassens et al., 2020; Dronsella et al., 2022) and *Pseudomonas putida* (Bruinsma et al., 2022; Turlin et al., 2022).

Despite the promising results on implementing synthetic formatotrophies, the engineered strains show too low growth-rates for industrially relevant bioproduction. Instead, natural formatotrophs could prove to be valid platforms for developing formate-based bioproduction. The facultative chemo-litho-autotroph *C. necator* is an ideal candidate to explore for this scope, as it is a capable of growing on formate as the sole carbon and energy source using the CBB cycle (Brigham, 2019; Grunwald et al., 2015; Li et al., 2012; Liu et al., 2016; Panich et al., 2021; Pavan et al., 2022; Raberg et al., 2018; Sohn et al., 2021; Volodina et al., 2016). This bacterium oxidizes formate to CO_2_ while reducing NAD^+^ to NADH through a kinetically fast (*k*_*cat*_ of 200 s^-1^) molybdenum-dependent formate dehydrogenase (Fdh) encoded by *fdsGBACD* (Niks et al., 2016).

The use of *C. necator* as cell factory has been facilitated thanks to a synthetic biology toolkit (Alagesan et al., 2018; Pan et al., 2021; Sydow et al., 2017) which, although limited, supported a plethora of metabolic engineering endeavors for the synthesis of value-added compounds. Historically, this bacterium has long been investigated for the endogenous synthesis of the biodegradable polymer poly(3-hydroxybutyrate) (PHB) (Li et al., 2020; Tang et al., 2020). Moreover, it has been engineered to convert fructose or, less frequently, CO_2_ and H_2_ into several other compounds such as, *e*.*g*., alka(e)ne (Crépin et al., 2016), the isoprenoids α-humulene (Krieg et al., 2018; Milker et al., 2021), β-farnesene (Milker and Holtmann, 2021), and lycopene (Wu et al., 2022), the alcohols isopropanol (Garrigues et al., 2020; Grousseau et al., 2014; Marc et al., 2017) and 1,3-butanediol (Gascoyne et al., 2021), acetoin (Härrer et al., 2021), methyl-ketones (Müller et al., 2013), as well as trehalose (Löwe et al., 2021), mannitol (Hanko et al., 2022), myo-inositol (Wang et al., 2023), and glucose (Wang et al., 2022).

These achievements certified the versatility of *C. necator* as cell factory for both heterotrophic and autotrophic cultivation modes. Yet, biological conversion of formate into value-added molecules has seldom been explored (*Table 1*). Apart from the endogenous polymer (PHB), the only platform chemicals derived from formate that are reported in literature are the alcohols isobutanol and 3-methyl-1-butanol (Li et al., 2012), the organic acids mesaconate, 2S-methylsuccinate (Hegner et al., 2020), and lactate (Kim et al., 2023). Most likely, the limited exploration of formate-based bioprocesses has a two-fold explanation: i) the lack of metabolic engineering endeavors targeted for formatotrophic bioproduction combined with ii) several cultivation challenges related to the use of formate as substrate. For the latter, several factors hamper formatotrophic growth. The first is the inherent low degree of reduction of the substrate, which requires high concentrations in the medium to support both microbial growth and product synthesis. Secondly, high concentrations of formate are toxic to most bacterial cells. Furthermore, as the deprotonated form of formate is mostly used for cultivations, its uptake inside the cell co-consumes protons, with consequent alkalinization of the medium which requires pH titration.

**Table 1.** Overview of added-value compounds obtained from biological conversion of formate.

| Organism | Product | Maximum biomass concentration | Product titers | Reference |
| --- | --- | --- | --- | --- |
| *Cupriavidus necator* | PHB | 0.193 g/L | 0.056-0.073 g/L | (Stöckl et al., 2020) |
| *Methylobacterium chloromethanicum* | PHB | ~ 5 g/L | 1.72 g/L | (Cho et al., 2016) |
| *C. necator* | PHB | 0.040 g/L | 0.031 g/L | (Janasch et al., 2022) |
| *C. necator* | PHB | 0.077 g/L | 0.005 g/L | (Jahn et al., 2021) |
| *C. necator* | PHB | < 1.0 OD<sub>600</sub> (0.363 g/L)<sup>a</sup> | 0.013-0.025 g/L | (Rowaihi et al., 2018) |
| *C. necator* | Isobutanol and 3-methyl-1-butanol (3MB) | ~ 4.0 OD<sub>600</sub> (1.5 g/L)<sup>a</sup> | ~0.846 g/L isobutanol and ~0.570 g/L 3MB | (Li et al., 2012) |
| *M. extorquens* | Mesaconate and 2S-methylsuccinate | ~ 0.14 OD<sub>600</sub> (0.05 g/L)<sup>a</sup> | 7 µM mesaconate and 10 µM of 2S-methylsuccinate | (Hegner et al., 2020) |
| *E. coli* | Lactate | ~ 1.53 OD<sub>600</sub> (0.56 g/L)<sup>a</sup> | 1.2 mM | (Kim et al., 2023) |
| *C. necator* | Crotonate | 7.8 OD<sub>600</sub> (2.83 g/L)<sup>a</sup> | 0.148 g/L or 1.74 mM | This study |

<sup>a</sup> the CDW value (expressed in g/L) was extrapolated using a CDW/OD<sub>600</sub> ratio of 0.363 (Grunwald et al., 2015).

Target compounds synthesized from formate should ideally serve as building blocks for the chemical industry. Organic acids are such a class of compounds, as they can be used as monomers for the synthesis of more complex molecules (Sauer et al., 2008; Becker et al., 2015). Unsaturated organic acids present an additional advantage due to their higher reactivity, which can be exploited for manufacturing. Therefore, exploring biological synthesis of unsaturated organic acids can offer new opportunities for producing building-blocks for a biobased economy.

The four-carbon molecule crotonate (also known as 2-butenoate, or trans-2-butenoate) is a short-unsaturated organic acid with biotechnological potential (Mamat et al., 2014). It can be used as building-block for the synthesis of several copolymers e.g., resins, surface coatings, plasticizers, and organic chemical intermediates (Li et al., 2021). However, its current production is petroleum-based (Wang et al., 2019), and only a handful of proof-of-concepts describe its microbial synthesis from heterotrophic substrates. These utilized the workhorse *E. coli* (Dellomonaco et al., 2011; Liu et al., 2015; Kim et al., 2016), and the emerging hosts *Methylobacterium extroquens* (Schada Von Borzyskowski et al., 2018) and *Yarrowia lipolytica* (Wang et al., 2019).

In this work, we targeted the two abovementioned challenges of formate-based bioprocesses and developed a proof-of-principle for the biological conversion of formate into crotonate using *C. necator* as cell factory. We started by assessing formate toxicity in the microorganism. This was followed by the implementation of a small-scale cultivation strategy using formate as only carbon and energy source, aiming at developing a setup for facilitating the screening of formatotrophic production strains. Then, we engineered a heterologous crotonate biosynthetic pathway in *C. necator*. To achieve this second objective, we used a modular approach by dividing the metabolic route into modules that were assessed both *via* enzymatic assays and *in vivo*. Finally, we harnessed the most promising pathway architecture in our cultivation strategy for demonstrating *in vivo* biological conversion of formate into crotonate in a continuous process.

## 2. Materials and methods

### 2.1 Bacterial strains, media, and standard cultivation conditions

The base strains and plasmids used in this work are listed in *Table 2*. LB medium (10 g/L bacto tryptone, 5 g/L yeast extract, 10 g/L NaCl) was used for standard cultivation of the strains *e*.*g*., for plasmid cloning, strain engineering, and strain preculturing. Antibiotics were used at the following concentrations: gentamycin, 20 µg/mL (native resistance of *C. necator*); kanamycin, 50 µg/mL (*E. coli*) or 100 µg/mL (*C. necator*). Moreover, 5-aminolevulinic acid was added at 50 µg/mL to support growth of *E. coli* ST18 conjugative strains (deleted in *hemA*).

**Table 2.**
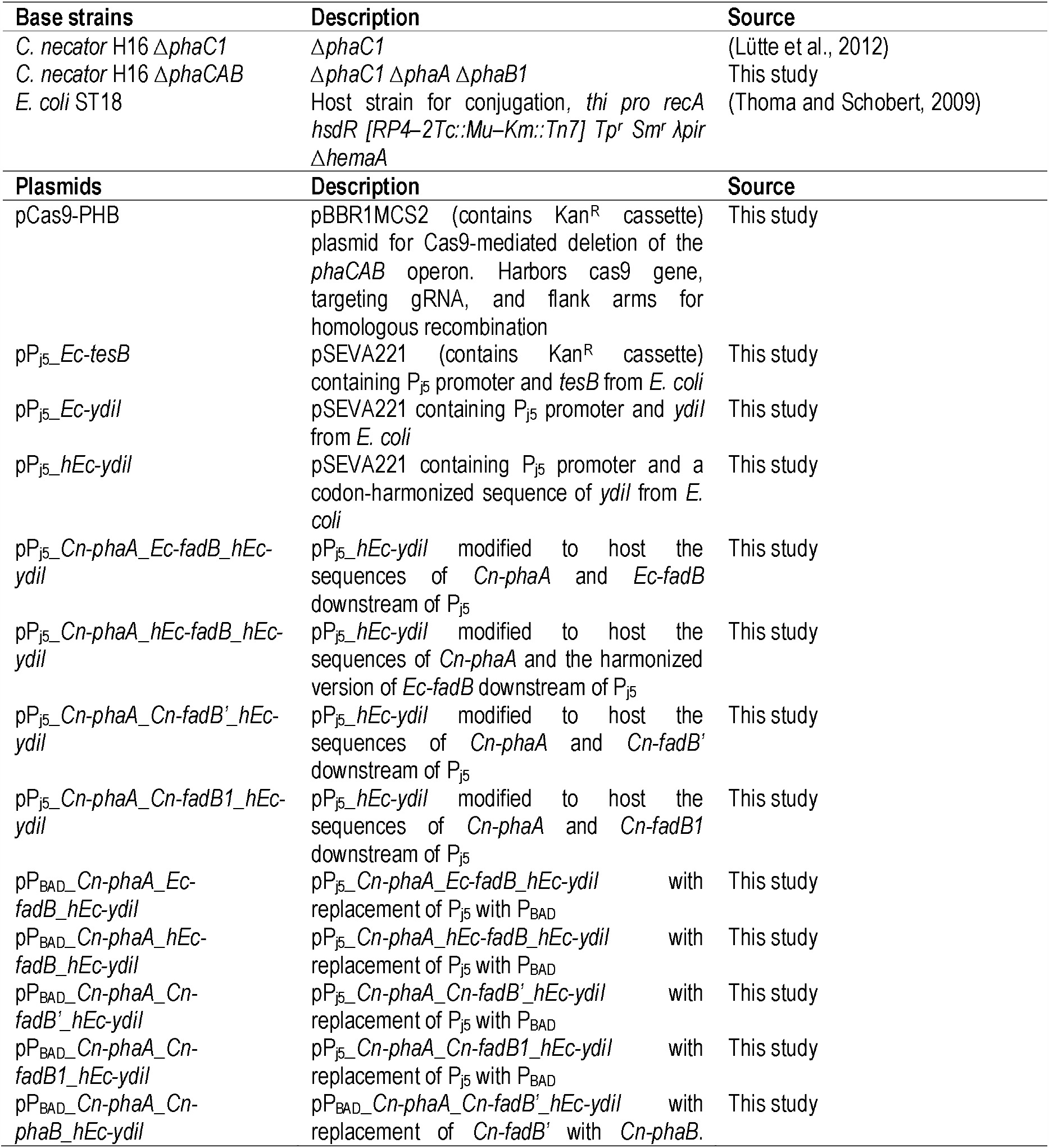
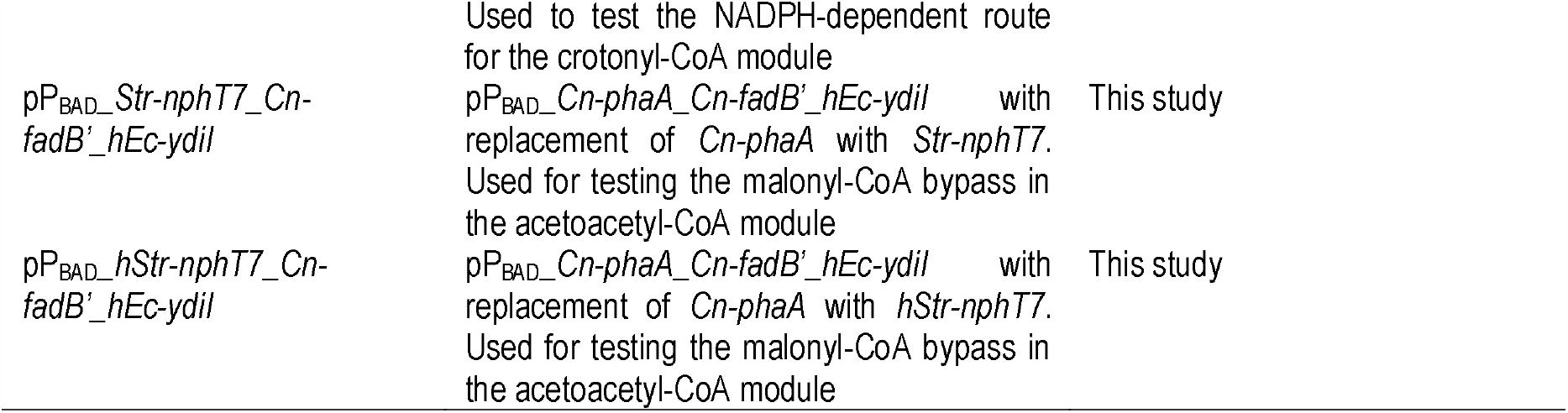
List of strains and plasmids used in this work.

### 2.2 Plasmid construction

Primers for PCR are listed in **Supplementary Table 1** and were synthesized by Integrated DNA Technologies (IDT). *C. necator* genomic DNA was extracted and used as template through the DNA purification kit from MACHEREY⍰NAGEL. Wild-type sequence of *Str-nphT7* (GenBank: AB272317) and codon harmonized gene sequences of *Ec*-*ydiI, Ec-fadB* and *Str*-*nphT7* (**Supplementary Table 2**) were ordered from Twist Bioscience. Codon harmonized sequences were generated using the Galaxy platform (Claassens et al., 2017). PCR amplifications were performed using Q5 High-Fidelity DNA polymerase Master Mix 2x from New England Biolabs (NEB). Due to the high GC content of *C. necator* genome, PCR reactions using its genomic DNA as template were supplemented with DMSO 3% v/v. All PCR products were purified via gel extraction using the Zymoclean Gel DNA Recovery kit (Zymo Research). Then, they were assembled employing the NEBuilder® HiFi DNA Assembly kit (NEB). The assembled plasmids were structured as follows: P_j5_ or P_BAD_ promoter (Alagesan et al., 2018; Gentz and Bujard, 1985), which controlled the expression of up to three genes, regulated in their translation by RBS1, RBS5, and RBS4 (Alagesan et al., 2018), respectively. Once constructed, plasmids were transformed into *E. coli* ST18 chemically competent cells, and plated on LB agar plates, supplemented with 50 µg/mL 5-aminolevulinic acid and 50 µg/mL kanamycin. Single colonies were then selected, grown overnight in liquid LB (supplemented with 50 µg/mL 5-aminolevulinic acid and 50 µg/mL kanamycin), and their plasmids were prepared using the GeneJET Plasmid Miniprep kit (Thermo Fisher Scientific). Once purified, plasmids were sent for Sanger sequencing to LGC Genomics.

### 2.3 Diparental conjugation of *C. necator*

Once the correct assembly was confirmed by sequencing, the corresponding *E. coli* ST18 strain was used as donor for diparental conjugation. Both *E. coli* ST18 and *C. necator* were precultured in liquid LB. This was supplemented or with 50 µg/mL 5-aminolevulinic acid and 50 µg/mL kanamycin (*E. coli* ST18), or gentamycin 20 µg/mL (*C. necator*). After overnight incubation, the strains were washed from the preculturing medium, and resuspended in LB (50 µg/mL 5-aminolevulinic acid) in a 3:1 ratio between *C. necator* and *E. coli*, respectively. Then, the combined strains were briefly centrifuged to collect their biomass as pellet, and this was resuspended in 50 µL, spotted on an LB agar plate supplemented with 50 µg/mL 5-aminolevulinic acid, and incubated at 30 °C for at least 5 hours. Then, the spot was resuspended in liquid LB supplemented with 100 µg/mL kanamycin and plated on LB agar plates supplemented with 100 µg/mL kanamycin. After 48-72 hours of incubation at 30 °C, single *C. necator* colonies were visible and selected for further testing.

### 2.4 Deletion of the PHB (*phaCAB*) operon in *C. necator*

For the deletion of the *phaCAB* operon, a pCas9 plasmid for homologous recombination and Cas9 counter-selection was cloned, following the approach previously described for *Rhodobacter sphaeroides* (Mougiakos et al., 2019). The spacers sp1 (5’- ACGCTTCCCGACCTACCGGA-3’) and sp2 (5’- ATGATGGAAGACCTGACACG- 3’) were cloned separately in two different pCas9 plasmids. 1 kb flanking arms for homologous were amplified from the genomic DNA of *C. necator*, corresponding to the regions upstream and downstream of the *phaCAB* operon. Deletion of *phaCAB* was obtained via diparental conjugation, and the primer set 6335/6336, annealing on the genomic DNA outside the flanking sites, was used for screening for deletions. Putative mutants were further checked with the primer set 6337/6338 binding internally within the *phaCAB* operon. After this second confirmation, the deletion was confirmed by Sanger sequencing.

### 2.5 Enzymatic assay for crotonyl-CoA thioesterase activity

Biological duplicates of *C. necator* strains harboring the pPj5 plasmids pP_j5_*_Ec-tesB*, pP_j5_*_Ec-ydiI*, and pP_j5_*_hEc-ydiI* were precultured overnight on liquid LB containing 20 µg/mL gentamycin and 100 µg/mL kanamycin. Then, they were washed in the same medium and inoculated with a starting OD_600_ of 0.1. After 3-5 hours, when the OD_600_ reached 0.5, the strain pellets were collected by centrifugation. The supernatant was removed, and the pellets were stored at -80 °C. Once all strains had been collected, pellets were thawed on ice, and membrane lysis was performed using the B-PER Reagent solution (Thermo Fisher). Each lysate was used for both enzyme assay and total protein quantification. The enzyme assay was performed as previously described (McMahon and Prather, 2014), using 5,5′-dithiobis-(2-nitrobenzoic acid), or DTNB, as colorimetric reagent for absorption detection at 412 nm. This molecule reacts in a 1:1 ratio with free CoA and, therefore, CoA can be quantified through the extinction coefficient for DTNB, which corresponds to 14.150 M^-1^ cm^-1^. The assay was performed in technical duplicates for each of the replicates in 96-well microplates (Nunclon Delta Surface, Thermo Scientific), using 150 µl of reaction volume, corresponding to a light path of approximatively 0.42 cm (https://static.thermoscientific.com/images/D20827~.pdf). Crotonyl-CoA was added as substrate at an initial concentration of 0.2 mM, in excess compared to DTNB (0.1 mM). To ensure a linear trend in the absorption during the assay, dilutions series of the cell lysates were performed. To quantify the total protein content of the lysates, we used the Lowry Assay kit from SERVA electrophoresis.

### 2.6 Heterotrophic cultivation in test tubes for crotonate production

Preculturing and cultivation of *C. necator* was performed in 4 mL of M9 medium (47.8 mM Na_2_HPO_4_, 22 mM KH_2_PO_4_, 8.6 mM NaCl, 18.7 mM NH_4_Cl, 2 mM MgSO_4_ and 100 μM CaCl_2_), supplemented with trace elements (134 μM EDTA, 31 μM FeCl_3_, 6.2 μM ZnCl_2_, 0.76 μM CuCl_2_, 0.42 μM CoCl_2_, 1.62 μM H_3_BO_3_, 0.081 μM MnCl_2_). As carbon source, 50 mM fructose was used. Additional 20 µg/mL of gentamycin and 100 µg/mL of kanamycin were added to the medium to select for plasmid-harboring *C. necator* strains. For the cultivation experiment, overnight grown precultures were washed in fresh 1X M9 salts media with no carbon source and diluted to an initial OD_600_ of 0.1 in 4 mL of fresh M9 medium. For cultivations requiring L-arabinose induction, growth was monitored by OD_600_ measurement and, once a value of 0.5 was reached, L-arabinose was supplemented in the media with the specific concentrations stated in the text. Cultivations were terminated after 24 hours from the initial inoculation, and the supernatant was collected by centrifugation for analytical analyses.

### 2.7 Microtiter plate cultivation for studying formate toxicity

For plate reader growth experiments, *C. necator* strains were revived from glycerol stocks and streaked on LB agar plates. Then, single colonies were picked and precultured in a sterile 15 mL test tube containing 4 mL of liquid LB overnight at 30°C and 250 rpm. Then, 40 µL of dense culture was used to inoculate a sterile test tube with 4 mL of M9 minimal medium supplemented with 80 mM formate or 20 mM fructose as carbon and energy source, followed by overnight incubation at 30°C and 250 rpm. Exponentially growing cell cultures were then washed two times in 1X M9 salts with no carbon source and used to inoculate the test medium (M9 medium supplemented with the indicated carbon sources in the text). The starting OD_600_ was set at 0.01; 150 µL of culture were added to each well and covered with 50 µL of mineral oil (Sigma-Aldrich) to prevent evaporation. The culture was incubated at 30°C in a microplate reader (EPOCH 2, BioTek) in the presence of 10% CO_2_ in the atmosphere. The shaking program cycle (controlled by Gen5 3.04) was performed as previously described (Wenk et al., 2020).

### 2.8 Formatotrophic cultivation in Erlenmeyer flask

Single *C. necator* colonies were precultured in 10 mL of LB broth, with 20 hours of incubation at 30 °C and 250 rpm. Then, cells were centrifuged and resuspended in JMM minimal medium (Li et al., 2012) containing: 64.0 mM Na_2_HPO_4_, 36.0 mM KH_2_PO_4_, 7.6 mM (NH_4_)_2_SO_4_, 0.8 mM MgSO_4_, 0.17 mM FeSO_4_, 0.02 mM CaCl_2_, 0.8 µM CoCl_2_, 0.5 µM MnCl_2_, 0.5 µM ZnCl_2_, 1 µM H_3_BO_3_, 0,1 µM Na_2_MoO_4_, 0.1 µM NiSO_4_, 0.1 µM CuSO_4_, 25 µM HCl, pH 6.8. As carbon sources, 55 mM fructose and 30 mM sodium formate were provided. After 6-8 hours, cultures were centrifuged, washed and transferred in 250 mL Erlenmeyer flasks containing 100 mL of fresh JMM medium supplemented with 5.4 g/L sodium formate (80 mM) as only carbon source. The starting OD_600_ was set at 0.1; the cultures were incubated at 30 °C and 250 rpm. When described, manual feeding for pH control was performed with 26 M formic acid solution (98%, Carl Roth) 3 times a day to bring the pH between 6.7 and 7.0.

### 2.9 Fed-batch formatotrophic cultivation in 2-L Minifors 2 bioreactors

Bacterial preculturing was performed by inoculating 10 mL of LB medium with fresh colonies of *C. necator* at 30°C and 250 rpm. After 6 hours of incubation, the cells were centrifuged and resuspended in 250 mL Erlenmeyer flasks (150 mL of working volume) containing JMM medium with 10 mM fructose and 50 mM of sodium formate as carbon sources. Throughout the preculturing, culture’s pH was kept between 6.7 and 7.0 by addition of 26 M formic acid. After 20 hours of incubation, the cultures were centrifuged and resuspended in 1L of fermentation mineral medium (Mozumder et al., 2014) which contained, per liter: 2 g (NH_4_)_2_SO_4_, 13.3 g KH_2_PO_4_, 1.2 g MgSO_4_•7H_2_O, 100 mg citric acid, and 10 mL trace element solution. The trace element solution contained: 10 g/L FeSO_4_•7 H_2_O, 2.25 g/L ZnSO_4_•7H_2_O, 1 g/L CuSO_4_•5 H_2_O, 0.5 gMnSO_4_•5 H_2_O, 2 g/L CaCl_2_•2 H_2_O, 0.23 g/L Na_2_B_4_O_7_•10 H_2_O, 0.1 g/L (NH_4_)6Mo_7_O_24_, and 35% HCl (10 mL/L). The carbon source (sodium formate), the nitrogen source (NH_4_)_2_SO_4_ and the magnesium sulfate (MgSO_4_•7H_2_O) were autoclaved separately and added afterwards. The culture medium was sparged with technical air and CO_2_ at a flowrate of 0.5 L/min. The partial pressure of CO_2_ and O_2_ in the exhaust gas were monitored with the CO_2_ and O_2_ BlueVary sensor (BlueSens GmbH). The partial pressure of CO_2_ (pCO_2_) was maintained at 5% by controlling the gas inflow. The dissolved oxygen concentration (pO_2_) was maintained above 30% of air saturation by controlling the agitation rate of the propeller from 500 rpm to 1600 rpm.

### 2.10 Fed-batch formatotrophic cultivation in 250-mL BlueLabs mini bioreactors

Inocula for fed-batch cultures were prepared in 150 mL of LB containing 100 µg/mL of kanamycin inoculated with a single colony of the tested strain. The cultures were incubated at 30°C overnight. The cultures were then resuspended in 100 mL of J Minimum Medium (abbreviated as JMM; 64.0 mM Na2HPO4, 36.0 mM KH2PO4, 7.6mM (NH4)2SO4, 0.8 mM MgSO4, 0.17 mM FeSO4, 0,02 mM CaCl2, 0.8µM CoCl2, 0.5 µM MnCl2, 0.5µM ZnCl2, 1µM H3BO3, 0,1µM Na2MoO4, 0.1 µM NiSO4, 0.1 µM CuSO4, 25µM HCl, pH 6.8) containing 55 mM fructose, 30 mM formate and 100 µg/mL of kanamycin. The cultures were incubated at 30°C, the pH regulated manually to 6.8-7.0 several times during the day by addition of 98% formic acid solution. After 8 hours of incubation, the cultures were centrifuged, and the pellet were resuspended in 2 times 150 mL JMM containing 80 mM formate and 100 µg/mL of kanamycin to reach an initial OD of 1.5. The strains were cultivated in 250 mL GLS80 flasks (Duran^®^) at 30°C. The cultures were stirred at 1000 RPM with a magnetic stirrer (diameter: 10 mm and length 60 mm) and the pH was controlled and monitored online by addition of 98% (26 M formic acid) with Bluelab pH Controller Connect (Bluelab). pH electrodes were decontaminated by 30 min incubation at 60°C in a solution with 0.5% bleach and rinsed abundantly with sterile culture medium. The production of crotonate was induced by adding 1 mM arabinose to the cultures after 17 hours incubation. Samples were taken regularly, and the supernatants were analyzed by HPLC.

### 2.11 Organic acid measurements with chromatography

Crotonate and 3-hydroxybutyrate (3HB) concentrations were measured via a Dionex ICS 6000 HPAEC Ion Chromatography (IC) system from Thermo Fischer, equipped with an organic acid column (Dionex IonPac AS11-HC-4µm) at 30 °C with a flow of 0.38 mL/min using ddH_2_O as eluent. To the eluent, KOH was added from a cartouche to create an alkaline fluid (pH 14), responsible of the ionization of the compounds and, therefore, their recognition in the conductivity detector. Quantification of the compounds was possible by creating calibration curves with known concentrations of crotonate and 3HB, which were prepared using commercially available standards of the molecules (Sigma-Aldrich). Concentrations of metabolites in the fed-batch samples were measured via the 1260 Infinity II LC system from Agilent equipped with an Hi-Plex 7.7 × 300 mm, 8 µm HPLC column (Agilent) at 60°C with a flow rate of 0.7 mL/min using 0.005 M H_2_SO_4_ in ddH_2_O as eluent. Metabolites were detected using a UV detector measuring at 210 nm and a refractive index detector at 55°C.

## 3. Results

### 3.1 Assessing formate toxicity in *C. necator* H16

Growth of *C. necator* on formic acid has been previously studied in batch (Lee et al., 2006), chemostat and a fed-batch setups (Grunwald et al., 2015). These works assessed formic acid toxicity for 0.5, 1.0, 2.0, and 5.0 g/L of substrate, corresponding to the range 10-110 mM assuming a molecular weight of formic acid of 46 g/mol.

As we were interested in achieving dense cell cultures, we hypothesized that it might be useful to assess the effect of formate toxicity by investigating also higher initial substrate concentrations. Therefore, we performed batch cultivations of a *C. necator* H16 strain deleted in its PHB biosynthetic pathway (Δ*phaCAB*) in a microtiter plate reader with sodium formate alone (formatotrophic setup) or in the presence of 20 mM fructose (mixotrophic setup) (*Fig. 1a*). In both cultivations, 10% CO_2_ was supplemented in the gas phase. We performed 1.5x serial dilutions of sodium formate from 1000 mM down to 17 mM (molecular weight of 68 g/mol). In both setups, formate concentrations above 197 mM (13.4 g/L) prevented growth. The highest final biomass concentrations reached on formatotrophic growth were in the range 59-132 mM of substrate (4.0-9.0 g/L), with final OD_600_ in the range 0.6-0.9 (*Fig. 1a*). When growing *C. necator* mixotrophically, we could detect a slight decrease in the final biomass concentration starting from 132 mM of formate (9.0 g/L), which became more evident at 198 mM (13.4 g/L) (*Fig. 1a*). The effect of formate toxicity was also confirmed when looking at the biomass yield on formate, which started to decrease for concentrations higher than 88 mM (*Fig. 1b*). When looking at the growth rates (expressed as doubling time (*Fig. 1c*), we observed that presence of formate in the mixotrophic cultivation had a negative effect on the doubling time, which was affected already at the lowest formate concentration tested (17 mM, corresponding to 1.2 g/L). Instead, in the formatotrophic setup the lowest doubling time was detected in the 39-88 mM (2.7-6-0 g/L) range.

**Figure 1.**
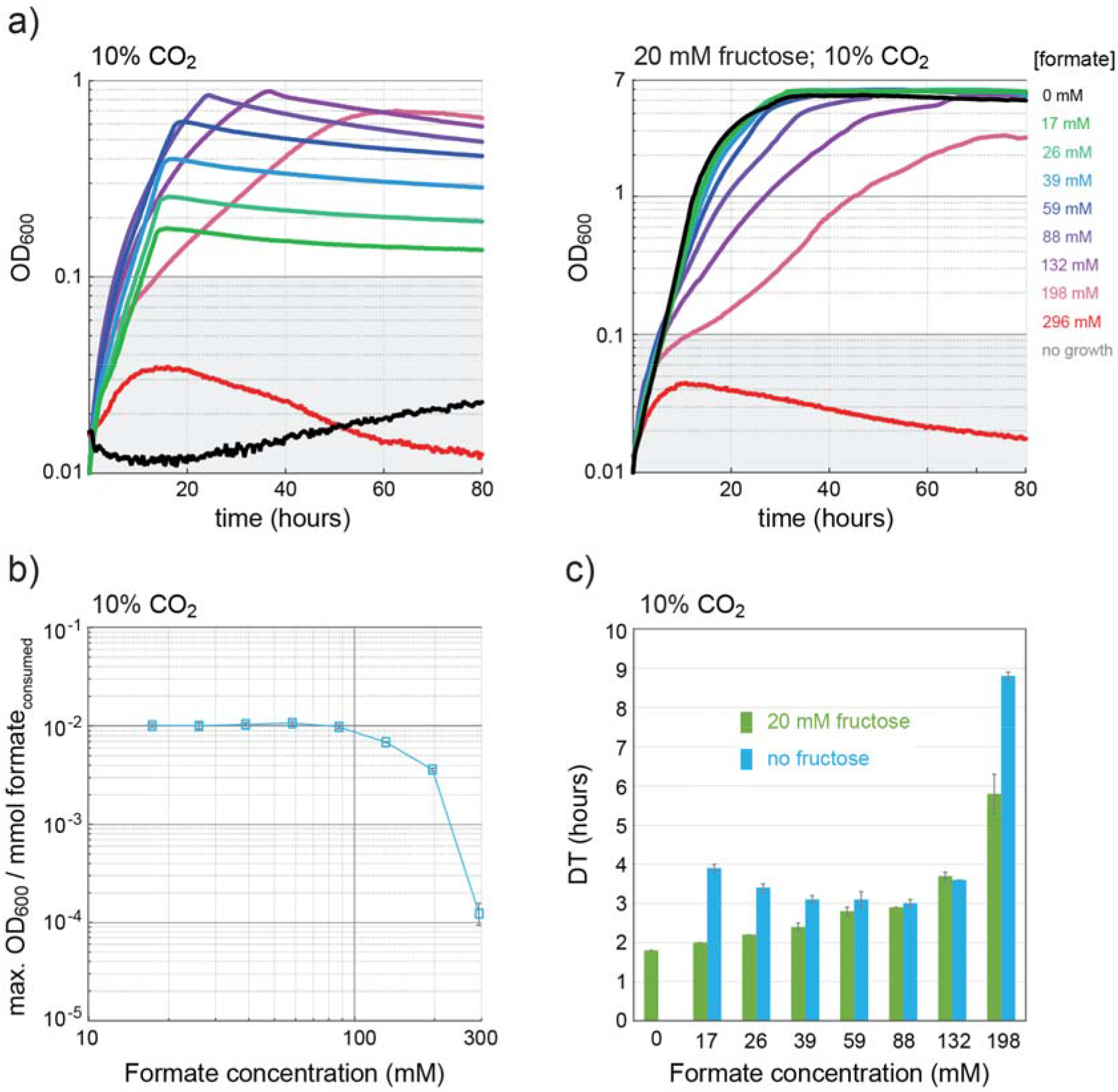
Formate toxicity assessment in *Cupriavidus necator* H16. a) A *C. necator* H16 strain (*ΔphaCAB*) was cultivated in a microtiter plate reader in the presence of different initial concentrations of formate. The effect of formate concentration on the growth profile was investigated for full formatotrophic growth mode (left) and mixotrophically in the presence of 20 mM fructose (right). b) Biomass yield on formate expressed as a function of the initial formate concentration in the medium. This relation has been calculated for the formatotrophic cultivation. c) Overview of the doubling time calculated for the formatotrophic and the heterotrophic growth experiments under different formate concentrations. The experiments were performed in biological triplicates and the error bars represent the standard deviation calculated with a 95% confidence interval.

This data confirms that a high concentration of formate (higher than 198 mM, 13.4 g/L) is incompatible with growth, independently from the presence of another substrate. When looking at formatotrophic cultivation, concentration of formate between 39-88 mM (2.7-6-0 g/L) resulted in the best tradeoff between growth rate and final biomass yields reached. We therefore used this information to guide the establishment of a system for the cultivation of *C. necator* in small-scale volumes.

### 3.2 Developing a small-scale formatotrophic cultivation setup for strain characterization

We decided to develop a small-scale cultivation setup compatible with *C. necator* growth, which can be later used to study product formation. We reasoned that such a setup can be used for the study of multiple strains, with the possibility of increasing the throughput for strain characterization. Our target was a cultivation strategy able to allow sufficient growth rates and cell densities using formate as only carbon and energy source.

To set a standard for comparison, we tested full formatotrophic growth in a 2-L benchtop bioreactor (1-L working volume) using a fed-batch setup with formic acid feeding. Here, we measured cell densities up to OD_600_ 23 after 50 h (**Fig. 2a**), corresponding to about 7.5-8.0 gCDW/L, and obtained a maximal growth rate of 0.29 h^-1^ during the initial batch phase.

**Figure 2.**
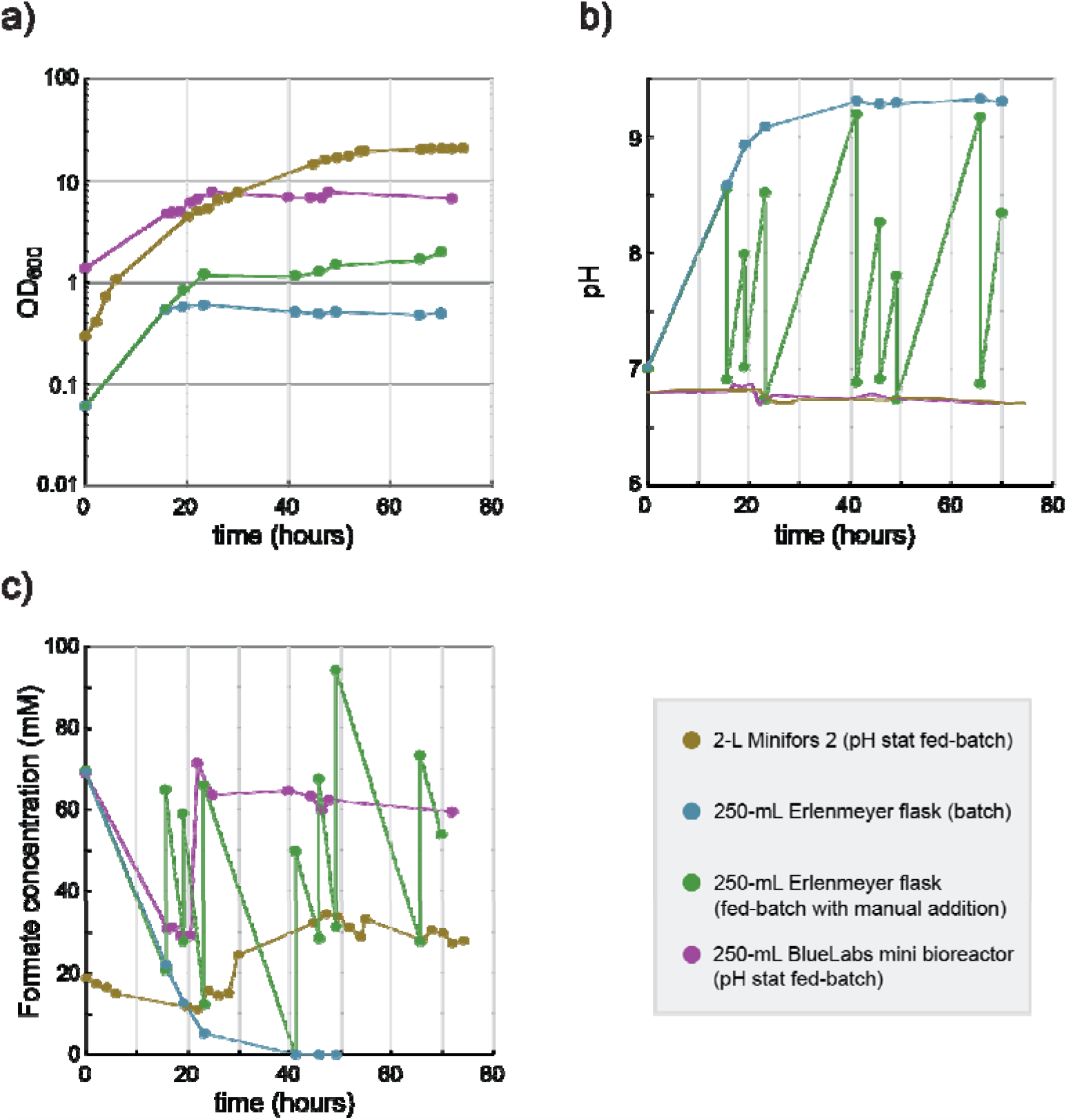
Implementation of a small-scale cultivation setup for formatotrophic growth of *C. necator* H16. a) Growth profile (OD_600_) of *C. necator* H16 under formatotrophic conditions. b) pH variation in the cultivation medium over the course of the cultivation. c) Variation in formate concentration in the medium over the course of the cultivation. The data shown represents the average of two biological duplicates for each cultivation setup tested.

Then, we focused on performing formatotrophic cultivations in a smaller setup, operating in the mL volume range. When studying formate toxicity, we observed that the optimal substrate concentration to support best growth rates and final ODs was within the 39-88 mM range (**Fig. 1**). We therefore performed a batch cultivation in 250-mL Erlenmeyer flasks (100 mL of cultivation volume) containing minimal medium supplemented with 70 mM formate. Here, *C. necator* growth resulted in a maximum OD_600_ of about 0.55 ± 0.00 (**Fig. 2a**). Moreover, formate consumption was associated with a strong increase of pH in the culture broth within the first 20 hours (**Fig. 2b**), reaching a value up to 9.3, which resulted in growth inhibition (**Fig. 2a, b**). Moreover, all the formate was depleted in the medium at 40 hours (**Fig. 2c**). Therefore, apart from its inherent toxicity, formate uptake by the cells resulted in rapid depletion from the broth with consequent medium alkalinization. To tackle these issues, we moved to a fed-batch setup where feeding of formic acid aimed at: i) maintaining the pH within viable range, ii) providing extra amounts of carbon and energy for biomass buildup, and iii) maintaining the substrate concentration within the non-toxic range of 39-88 mM.

We approached the small-scale fed-batch cultivation by first attempting manual feeding of formic acid to 250-mL shake flask cultures (100 mL working volume) supplemented with an initial formate concentration of 70 mM. Here, formic acid feeding allowed to extend both the cell viability and growth phase while increasing the final biomass concentration almost four-fold, reaching a maximum OD_600_ of 1.98 ± 0.04 within 70 hours (**Fig. 2a**). However, in this setup the growth rate is quickly limited by the frequency of manual feedings. In addition, the culture is alternating from abundance to depletion of formate (**Fig. 2c**), with consequent fluctuations on the medium pH (**Fig. 2b**). Therefore, the manual feeding strategy is not suitable to study optimal production conditions. We therefore reasoned to move to an alternative system where pH is measured online and kept constant by pump-feeding (pH stat). Such an approach allowed us to control the pH in 250-mL mini-stirred tank bioreactors, using 150 mL of working volume. This strategy significantly improved the cultivation performances, enabling us to maintain the pH around 6.8 while supplying continuously formic acid to the cultures (**Fig. 2 a-c**). Since we could automatically counter-balance medium alkalinization, we performed an inoculum with a higher starting OD_600_ > 1.0 (**Fig. 2a**), aiming at reaching high biomass concentration in a shorter time. Indeed, within the first 24 hours we obtained OD_600_ up to 7.80 ± 0.45, corresponding to 2.7-3.0 gCDW/L, and growth rates between 0.08 and 0.10 h_^-1^_.

Overall, this setup resulted in a 15-fold improvement in maximum biomass concentrations compared to the batch cultivation in flasks. Moreover, it allowed us to obtain cell densities within the g/L range within 24 hours of cultivation. We concluded that this setup succeeded in growing *C. necator* on formate at a satisfactory growth-rate and -yield for exploring formate-based bioproduction in engineered strains.

### 3.3 Metabolic engineering of *C. necator* for implementing a crotonate biosynthetic pathway

#### 3.3.1 Dividing crotonate biosynthesis into functional modules

After developing a cultivation setup for formatotrophic growth, we focused on establishing *C. necator* as cell factory to produce crotonate. We first identified possible enzymatic reactions that can lead to crotonate production. These biosynthetic routes branch from central carbon metabolism at the level of acetyl-CoA. We divided the pathway into three functional modules, each named after the product molecule of the respective module (**Fig. 3**).

**Figure 3.**
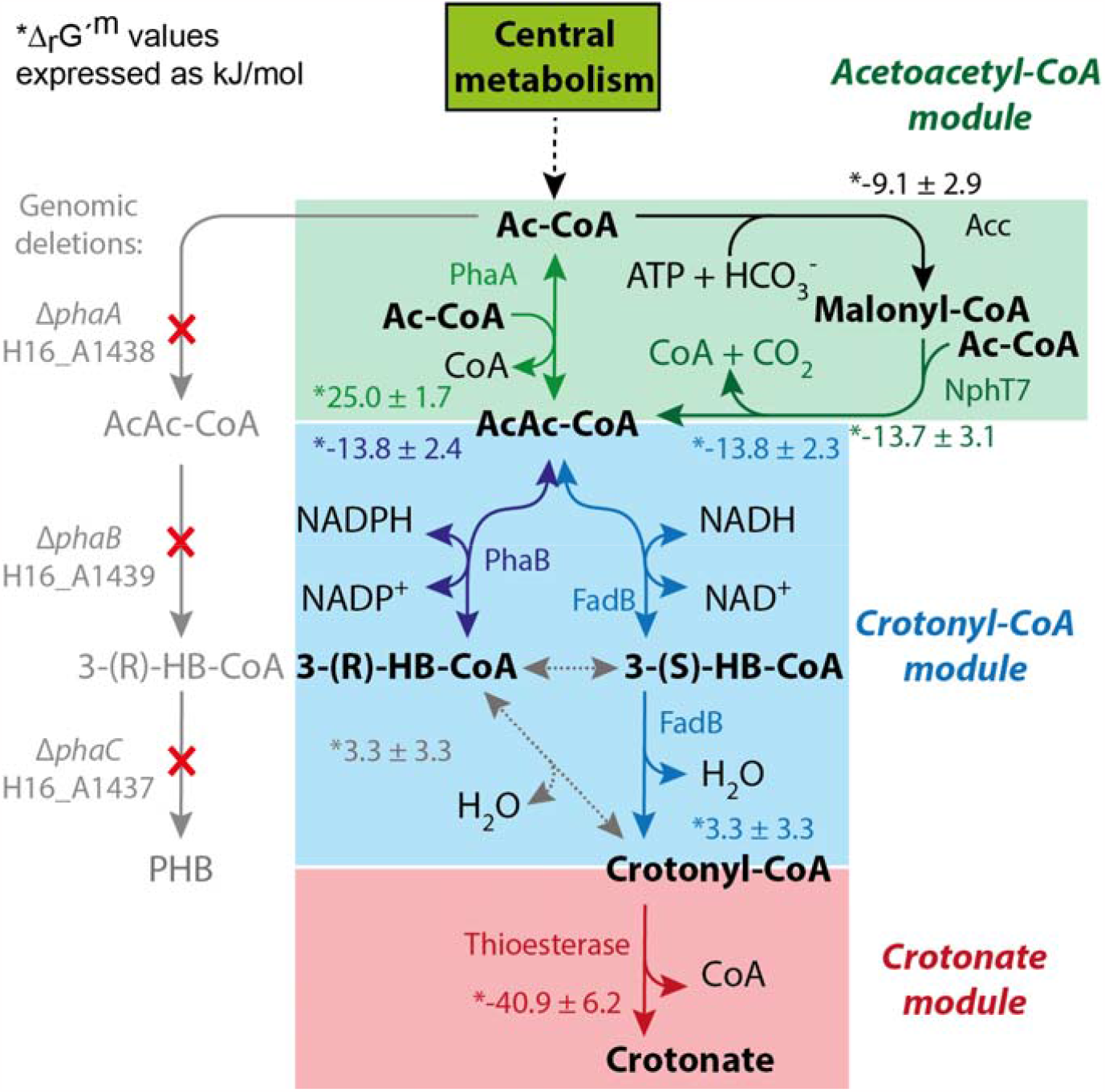
Overview of the engineered metabolic network described in this work leading to crotonate biosynthesis in *Cupriavidus necator*. The network is divided into three modules, each defined by its product. The first module (green box) is responsible for the synthesis of acetoacetyl-CoA. We assessed two possible routes, one mediated by PhaA, and the other via a malonyl-CoA dependent bypass (Acc and NphT7). The second module (light blue box) reduces and subsequently dehydrates acetoacetyl-CoA to crotonyl-CoA. We envisioned two possible routes for this module, one going through NADPH-mediated reduction via PhaB (overlapping with PHB biosynthesis), and the other occurring via NADH-mediated reduction (through reverse β-oxidation, FadB). The last module converting crotonyl-CoA to crotonate was investigated by testing different thioesterase candidates. For each reaction investigated, we included the estimated Δ_r_G′^m^. To prevent interference of the endogenous metabolism to product formation, the phaCAB operon (H16_A1437-A1439) was deleted. Abbreviations: acetyl-CoA (Ac-CoA), acetoacetyl-CoA (AcAc-CoA), 3-(R)-hydroxybutyryl-CoA (3-(R)-HB-CoA), 3-(S)- hydroxybutyryl-CoA (3-(S)-HB-CoA).

Subsequently, we explored the metabolic context of *C. necator* through the Kyoto Encyclopedia of Genes and Genomes database (KEGG) (Kanehisa and Goto, 2000) to identify endogenous enzymes for each module. To include other enzyme candidates in our study, we additionally included at least one non-native enzyme for each module, which had been previously characterized in other hosts.

The first section of the pathway, the ‘acetoacetyl-CoA module’, connects acetyl-CoA to acetoacetyl-CoA, and can be achieved via two possible routes. The first one consists of a single enzymatic step, catalyzed by the thiolase acetyl-CoA *C*-acetyltransferase (EC 2.3.1.9), encoded by the endogenous gene *phaA*, which condenses two acetyl-CoA molecules. The alternative route is supported by two reactions, the first one involving the ATP-dependent carboxylation of acetyl-CoA to malonyl-CoA (catalyzed by acetyl-CoA carboxylase, EC 2.1.3.15), encoded by endogenous *accABCD*. Then, malonyl-CoA is condensed with another acetyl-CoA via a decarboxylative Claisen condensation catalyzed by acetoacetyl-CoA synthase (EC 2.3.1.194). This last step requires the heterologous expression of *nphT7*, a gene originating from *Streptomyces* sp. *CL190* (Okamura et al., 2010). In addition to testing the wildtype coding sequence of *nphT7* (*Str-nphT7*), we additionally tested a codon-harmonized *hStr-nphT7* sequence (Angov et al., 2011; Claassens et al., 2017), which matches the codon usage of *C. necator*.

The following ‘crotonyl-CoA module’ is responsible for the conversion of acetoacetyl-CoA to crotonyl-CoA. This module catalyzes a reduction to 3-hydroxybutyryl-CoA (3-HB-CoA), followed by a dehydration step to crotonyl-CoA. In principle, two possible routes can be explored for this module in *C. necator*. The first one consists of a section of the reverse β-oxidation pathway (Dellomonaco et al., 2011). It involves a NADH-dependent reduction of acetoacetyl-CoA, generating the 3-(*S*)-HB-CoA enantiomer, and is followed by a dehydration step. One enzyme, FadB, contains the two catalytic domains required, and was shown to support crotonate synthesis in engineered *E. coli* (Kim et al., 2016). We therefore selected the *E. coli fadB* coding sequence (*Ec-fadB*) and its codon harmonized sequence (*hEc-fadB*) as candidates. Moreover, we explored possible endogenous *C. necator* candidates, as this bacterium encodes three *fadB* variants (named *Cn-fadB1* (H16_A1526), *Cn-fadB2* (H16_B0724), and *Cn-fadB’* (H16_A0461) (Insomphun et al., 2014). As *Cn-fadB1* and *Cn-fadB’* were shown to be the most catalytically active variants (Brigham et al., 2010; Volodina and Steinbüchel, 2014), we included them among the candidates to test.

The second option for the ‘crotonyl-CoA module’ partially overlaps with the PHB biosynthetic route. PhaB, which catalyzes the NADPH-dependent reduction of acetoacetyl-CoA to 3-(*R*)-HB-CoA, is the responsible enzyme for this first conversion. Three homolog genes *phaB1, phaB2* and *phaB3* exist in *C. necator*, of which *phaB1* is the main isozyme responsible for PHB biosynthesis (Budde et al., 2010). Further conversion of 3-(*R*)-HB-CoA to crotonyl-CoA requires an (*R*)-stereospecific dehydratase step, catalyzed by an (*R*)-specific enoyl-CoA hydratase (EC 4.2.1.119), such as PhaJ. *C. necator* is not known to produce this enzyme, although several *phaJ* homolog genes exist in this bacterium (Kawashima et al., 2012).

Finally, the ‘crotonate module’ catalyzes the thioesterase activity removing the CoA moiety from crotonyl-CoA, generating crotonate as product. YdiI catalyzes this reaction in *E. coli* (Kim et al., 2016), albeit with a low specific activity (McMahon and Prather, 2014). We selected *Ec-ydiI* and its codon-harmonized *hEc-ydiI* version to test in *C. necator*. Other enzymes, such as TesB, are known to be fast thioesterases while presenting a broad substrate range (McMahon and Prather, 2014). After having identified all enzyme variants, we investigated each module separately.

#### 3.2.2 Screening of crotonyl-CoA thioesterase candidates for the crotonate module

As no specific crotonyl-CoA thioesterase enzymes are known, we started our screening focusing on the last module of the pathway. We decided to first determine the activity of interest using cell lysates. Therefore, we cloned the genes of *Ec-ydiI*, its codon harmonized version (*hEc-ydiI*), and the broad-range thioesterase (*Ec-tesB*) gene each under the control of the strong constitutive promoter P_j5_ (Gentz and Bujard, 1985). Then, we cloned them separately into a pSEVA221 plasmid (RK2 origin of replication). We conjugated these plasmids in *C. necator* Δ*phaC1*, and performed an *in vitro* assay as previously described (McMahon and Prather, 2014), relying on the colorimetric signal resulting from the conversion of DTNB to TNB monitoring the release of free CoA (Fig. 3a).

The assay, which is based on the interaction of free CoA with DTNB to generate TNB, was initiated with addition of 0.2 mM of crotonyl-CoA to cell lysates (**Fig. 4a, Supplementary Fig. S1**). The TNBs signal from this assay suggested that Ec-TesB supported the highest reaction rate towards crotonate, followed by the codon harmonized version of YdiI (hEc-YdiI) (**Fig. 4b**). Nevertheless, a latent thioesterase activity was determined also in the negative control, which contained an empty pSEVA221 (**Fig. 4b**). This result was likely due to the reaction of intracellular free CoA, already present in the lysates, with DTNB.

**Figure 4.**
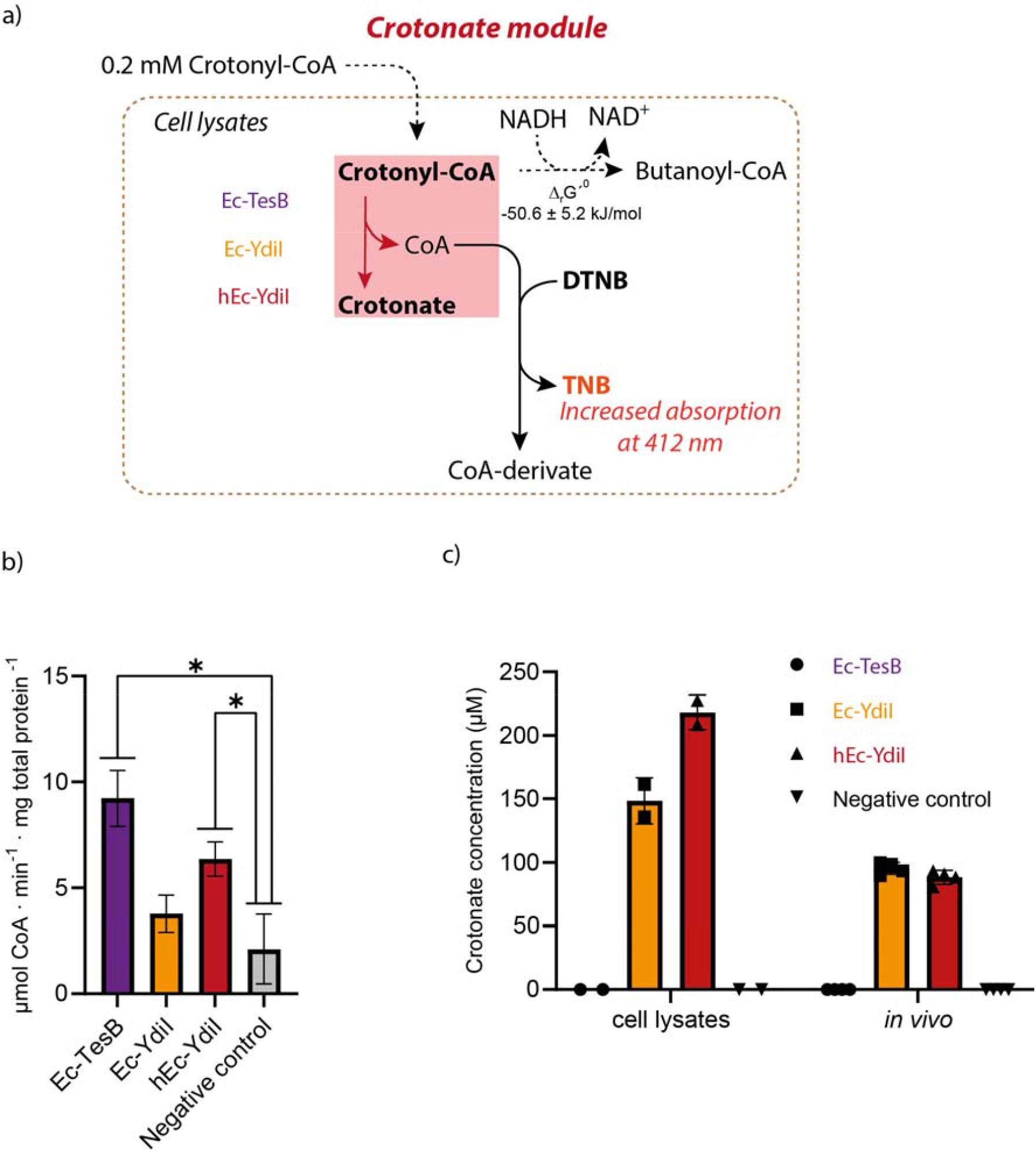
Assessment of the thioesterase activities for the crotonate module. a) Schematic of the enzyme assay mechanism using (5,5′-dithiobis-(2-nitrobenzoic acid) (DTNB) as colorimetric reactant on cell lysates. b) *V*_*MAX*_ of the different enzyme determined in cell lysates expressing the candidate enzymes. The values were determined through analysis of biological duplicates, each one analyzed in triplicate. The error bars represent the standard deviation between replicates. The asterisk indicates significant difference to the negative control (*p*-value < 0.05). c) Crotonate measured in the reaction wells after the enzymatic assay (cell lysates), as well as in the spent medium after 24 h incubation (*in vivo*, with fructose as substrate). Each data point represents a measurement from a biological replicate. For the cell lysates, we measured two biological replicates, whereas for the *in vivo* analysis, we measured crotonate on four biological replicates.

To overcome this experimental limitation, we measured the crotonate concentration in the spent assay medium *via* ion-chromatography (IC). To our surprise, Ec-TesB did not perform any conversion of crotonyl-CoA into crotonate. As the network around crotonyl-CoA allows other reactions to occur (**Fig. 4a**), we speculate that Ec-TesB might have contributed to the release of CoA from other metabolites that were present in the lysate or generated by enzymes in the lysate from the crotonyl-CoA supplemented at the beginning of the assay. However, in contrast to Ec-TesB, both YdiI variants clearly converted crotonyl-CoA into crotonate (**Fig. 4c**).

Ultimately, we assessed the ability of these three enzymes to support crotonate biosynthesis *in vivo* under heterotrophic fructose cultivation relying on native pathway enzymes for the upstream activities. We selected fructose as substrate as *C. necator* is unable to catabolize glucose naturally (Volodina et al., 2016). Indeed, we could confirm the exclusive ability of both YdiI variants to support similar levels of crotonate biosynthesis around 100 µM (**Fig. 4c**). Due to the slightly higher *V*_MAX_ of the harmonized version of YdiI in the *in vitro* assay (**Fig. 4b**), we selected hEc-YdiI as best candidate for the crotonate module.

#### 3.3.3 Studying the crotonyl-CoA module

##### 3.3.3.1 Choice of *fadB* homologs for the NADH route

Next, we studied the conversion of acetoacetyl-CoA into crotonyl-CoA. As native PHB biosynthesis can in principle compete with this module by consuming acetoacetyl-CoA - the *K*_M_ of PhaB for acetoaceyl-CoA is 5.7 µM (Matsumoto et al., 2013) - we deleted the whole *phaCAB* operon from the *C. necator* genome. To generate this Δ*phaCAB* strain, we used the CRISPR/Cas9 method previously described for *R. sphaeroides* (Mougiakos et al., 2019), which is structurally similar to the previously described toolbox for *C. necator* (Xiong et al., 2018). Although a knock-out mutant was detected and the deletion was confirmed by sequencing (**Supplementary Fig. S2**), the editing efficiency was lower than 10%, probably because the synthetic promoter for the gRNA transcript was specific for *R. sphaeroides* (Huo, 2011). Nevertheless, the successfully generated Δ*phaCAB* strain could be used as a platform for testing enzyme candidates.

As mentioned above, two key reactions make up this module: reduction of acetoacetyl-CoA to 3-hydroxybutyryl-CoA (3-HB-CoA), and dehydration of 3-HB-CoA to crotonyl-CoA (**Fig. 5a**). In principle, FadB alone (overlapping with reverse β-oxidation, NADH route) can support both activities, as the *fadB* gene from *E. coli* (*Ec-fadB*) has been already described to support crotonate biosynthesis (Kim et al., 2016).

**Figure 5.**
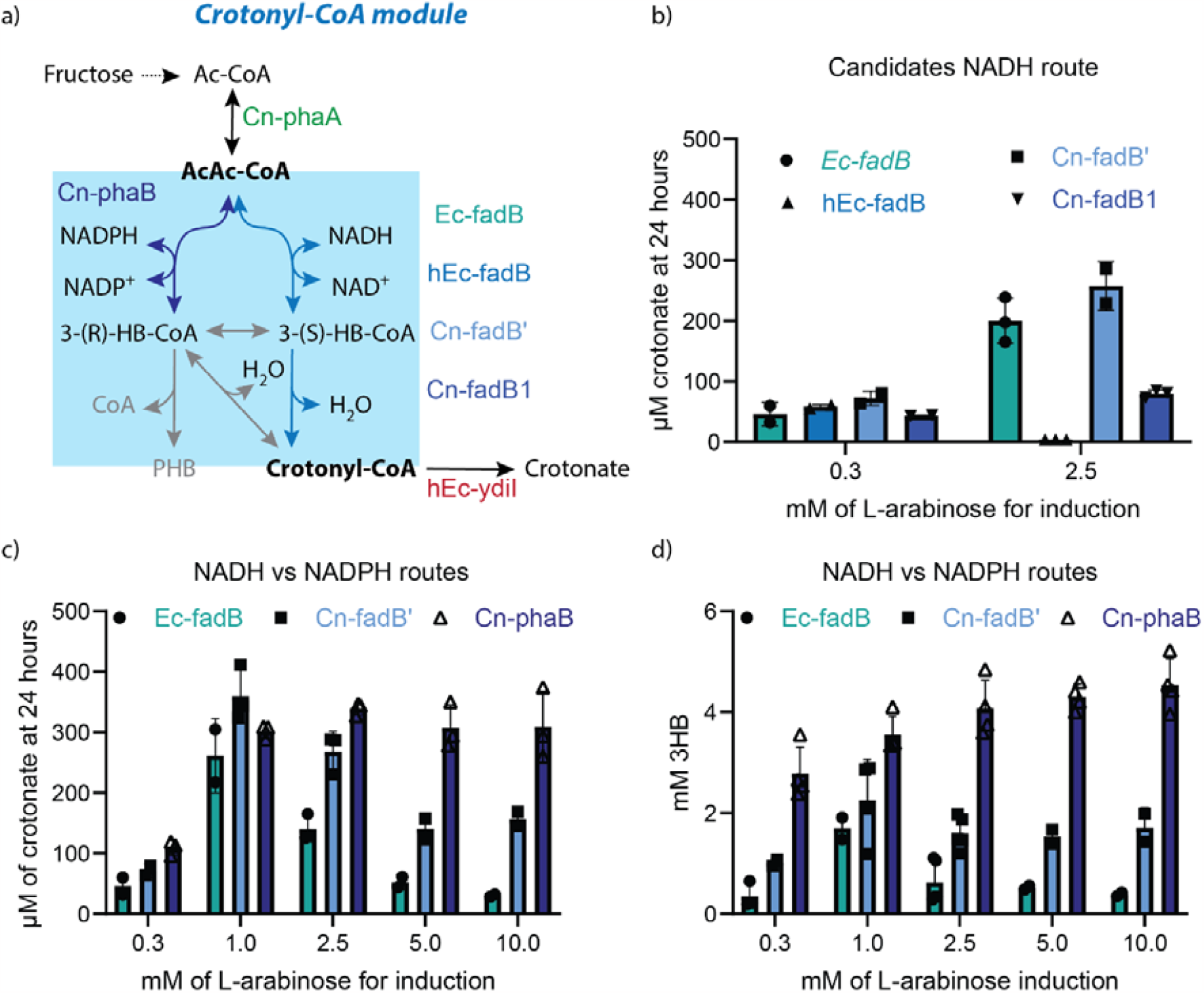
Study of candidates for the crotonyl-CoA module *via* NADH- and NADPH-dependent routes. a) Overview of the network supporting conversion of acetoacetyl-CoA (AcAc-CoA) into crotonyl-CoA. To aid in the screening, the *phaCAB* operon was deleted. Also, *Cn-phaA* and *hEc-ydiI* were expressed together with the module candidates on a plasmid. Cultivations were performed using fructose as substrate. b) Test of FadB orthologs as candidates for the NADH-dependent route. The assessment was performed using an L-arabinose inducible promoter, and crotonate was measured in the spent medium after 24 hours incubation. c, d) Characterization of the Ec-FadB and Cn-FadB’ orthologs (NADH route) and Cn-PhaB (NADPH route) over a wider range of induction with L-arabinose after 24 hours of cultivation. Panel c) reports crotonate titers, whereas panel d) shows the secretion of 3-(R/S)-hydroxybutyrate (3HB) in the spent medium. Each data point represents the measurement of a biological replicate. The bars indicate the standard deviation calculated on a dataset involving biological duplicates or triplicates.

The candidate genes for this module were cloned individually upstream of *hEc-ydiI* in the pSEVA221 plasmid. To reestablish the connection from acetyl-CoA in the Δ*phaCAB* strain, we also included *Cn-phaA* as first gene of the operon, directly under the control of the P_j5_ promoter. We also used medium-strength RBSs upstream of each gene to ensure good expression levels (Alagesan et al., 2018).

We then performed cultivations on fructose to determine the best candidate for the crotonyl-CoA module. To our surprise, no crotonate was detected in the spent medium of the different strains after 24 hours, except for about 80 µM of crotonate for the construct containing *Cn-fadB’*. As all constructs contained *hEc-ydiI*, which we had previously shown to support crotonate production *in vivo*, we hypothesized an instability of the engineered plasmids due to toxic levels of gene expression via the constitutive P_j5_ promoter. Therefore, at the end of the cultivation, we sequenced the crotonate biosynthetic plasmids from these strains. Indeed, we could observe multiple mutations or missing gene parts in all the constructs (data not shown), which suggested toxic expression levels.

To ensure more stable and controllable gene expression, we replaced in all constructs the constitutive P_j5_ promoter with an L-arabinose inducible P_BAD_ promoter. We selected this induction system as it displays a rather linear and wide dynamic range of expression in *C. necator* (Alagesan et al., 2018). Moreover, L-arabinose is not catabolized by *C. necator*, and therefore would not be used as carbon source (Alagesan et al., 2018; Pan et al., 2021). As the expression range of P_BAD_ has already been characterized in *C. necator* (Alagesan et al., 2018), we tested the values corresponding to approximately 50% and 100% induction levels using a fluorescent report protein (0.3 and 2.5 mM of L-arabinose, respectively) to screen the different FadB candidates. While the difference between the four candidates was limited at 0.3 mM, at 2.5 mM Ec-FadB and Cn-FadB’ clearly performed best, reaching crotonate titers of at least 200 µM (**Fig. 5b**). We speculate that in the *hEc-fadB* construct the toxicity issue of high expression levels might have persisted, as induction with 0.3 mM L-arabinose resulted in crotonate synthesis, whereas induction with 2.5 mM L-arabinose did not (**Fig. 5b**).

##### 3.3.3.4 Comparing NADH and NADPH routes to crotonyl-CoA

We next compared crotonyl-CoA formation through NADH- (partial reverse β-oxidation) to the NADPH-dependent route. The latter requires the native PhaB1 (Cn-PhaB), which is a well-characterized enzyme for 3- (*R*)-HB-CoA production in canonical PHB synthesis. However, Cn-PhaB is not expected to catalyze the further dehydration step to crotonyl-CoA. Therefore, endogenous reactions would be needed, either for the isomerization of 3-(*R*)-HB-CoA to 3-(*S*)-HB-CoA, or for the dehydration of 3-(*R*)-HB-CoA to crotonyl-CoA. For the latter reaction, 16 putative *R*-enoyl-CoA dehydratases have been identified in *C. necator* (Kawashima et al., 2012). Therefore, we opted to explore if, upon *Cn*-*phaB1* overexpression, an endogenous dehydratase activity could support *in vivo* crotonyl-CoA synthesis.

To properly compare the NADH- and NADPH-routes, we cultivated the strains harboring *Ec-fadB, Cn-fadB’*, and *Cn-phaB* with a wider range of L-arabinose induction, ranging from 0.3 to 10.0 mM, and measured crotonate titers after 24 h (**Fig. 5c**). We observed that, for the two FadB orthologs, the optimal induction level centered around 1.0 mM L-arabinose, with Ec-FadB producing about 250 µM crotonate, and Cn-FadB’ about 350 µM (**Fig. 5c**). Overall, Cn-FadB’ performed constantly better than Ec-FadB, supporting crotonate titers above 100 µM also at 5.0 and 10.0 mM of L-arabinose induction, whereas the decrease in titers for Ec-FadB was sharper (**Fig. 5c**). Instead, the Cn-PhaB strain resulted in a different behavior, with crotonate titers reaching a plateau of 300 µM in all replicates, except for the lowest induction level of 0.3 mM L-arabinose, which reached a lower concentration of 100 µM crotonate (**Fig. 5c**).

As PHB biosynthesis is inactivated in the Δ*phaCAB* strain, we anticipated to observe secretion of metabolic intermediates in the medium. In particular, we looked at 3-hydroxybutyrate (3HB) secretion by measuring the cumulative 3-(*R*)-HB and 3-(*S*)-HB concentrations in the spent medium by IC (**Fig. 5d**). The profile of extracellular 3HB (**Fig. 5d**) resembles the one of crotonate generated by the different strains (**Fig. 5c**), although with titers higher by an order of magnitude (mM range).

For the strains harboring the FadB enzymes, intracellular accumulation and consequent secretion of 3HB is most likely due to the kinetic properties of the following YdiI. This enzyme, presenting a *K*_M_ for crotonyl-CoA of 524 μM, and *k*_cat_ of 0.93 s^-1^ (Kim et al., 2016), is kinetically suboptimal. In fact, when compared to the average kinetic values of metabolic enzymes (Bar-Even et al., 2011), YdiI displays about 5-fold higher *K*_M_ and a 10-fold lower *k*_cat_ than the respective median values of 100 µM (*K*_M_) and 10 s^-1^ (*k*_cat_). Therefore, YdiI might not consume crotonyl-CoA at the same rate as it is produced by FadB, leading to its accumulation as well as 3-(*S*)-HB upstream of the pathway triggering 3HB secretion. Moreover, we speculate that the decreased titers of crotonate and 3HB measured at high induction levels for the two FadB enzymes (2.5 to 10.0 mM L-arabinose) are likely related the toxic effect of their gene expression.

The PhaB harboring strain regularly secreted higher amounts of 3HB compared to the FadB strains (*Fig. 5d*). At least two complementary interpretations are possible for this behavior. The first one is that 3-(*R*)-HB-CoA is produced and further converted to crotonyl-CoA by the endogenous (*R)*-enoyl-CoA dehydratases. However, the latter endogenous activity is not efficient enough, and 3-(*R*)-HB-CoA accumulates, with consequent secretion of 3HB. The second interpretation would consider 3-(*R*)-HB-CoA as a dead-end metabolite, which is eventually secreted, and production of crotonyl-CoA is supported by the endogenous metabolism, as for the case of *in vivo* production of crotonate via the P_j5__*hEc-ydiI* construct (**Fig. 2c**). Both interpretations are coherent with what was recently reported in literature, where the (*S*)-, and not (*R*)-, stereospecificity of enoyl-CoA hydratase activity is the most dominant in *C. necator* (Segawa et al., 2019).

Eventually, the NADH-route supported by Cn-FadB’ resulted in higher crotonate titers and lower secretion of 3HB compared to the NADPH-route. Hence, we chose to continue using the NADH-route as candidate for the crotonyl-CoA module.

#### 3.2.4 Engineering the acetoacetyl-CoA module using a thermodynamically favorable bypass

The enzyme acetyl-CoA *C*-acetyltransferase (PhaA), which condenses two acetyl-CoA into acetoacetyl-CoA, has an inherent thermodynamic limitation. In fact, the Δ_r_G^′m^ calculated for this reaction is 26.1 ± 1.7 kJ/mol (Flamholz et al., 2012). When growing on sugars, this thermodynamic barrier is lowered by a high intracellular level of acetyl-CoA, which is caused by nutrient (*e*.*g*., nitrogen) limitation in the medium (van Wegen et al., 2001). When growing on a C1 feedstock (*e*.*g*., formate or CO_2_), this scenario is unlikely to occur. In fact, during growth on formate, flux towards acetyl-CoA is reduced in *C. necator* (Jahn et al., 2021). A possible interpretation is that formate oxidation by formate dehydrogenase (Fdh) provides enough NADH to the cell, and the TCA cycle does not have to operate for energy generation.

We therefore opted to engineer an ATP-driven bypass of the PhaA reaction (linear pathway), aiming at increasing the thermodynamic drive for acetoacetyl-CoA generation (**Fig. 6a**). This new reaction is based on the combined activity of acetyl-CoA carboxylase (Acc) and NphT7, an acetoacetyl-CoA synthase which catalyzes the decarboxylative condensation of malonyl-CoA to acetyl-CoA to generate acetoacetyl-CoA (Okamura et al., 2010). The combination of ATP hydrolysis, bicarbonate fixation and direct decarboxylation together make this route thermodynamically far superior to the linear PhaA dependent route. The use of NphT7 in combination with Acc already demonstrated to support production of isobutanol (Lan and Liao, 2012) and 3-hydroxybutyrate (Ku and Lan, 2018) in the cyanobacterium *Synechococcus elongatus*. Whereas in *C. necator* the production of NphT7 requires heterologous gene expression, the Acc complex is endogenous and represents the first step of fatty acid biosynthesis.

**Figure 6.**
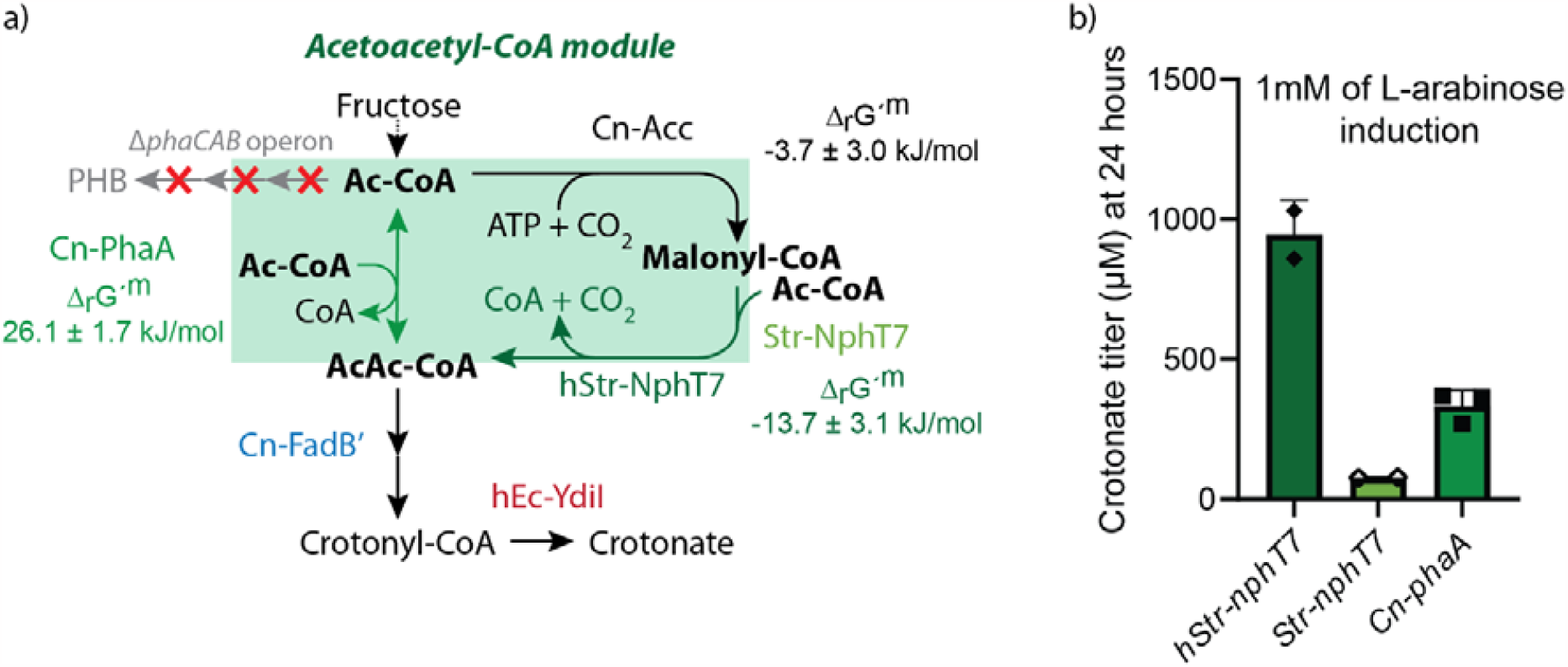
Condensation of acetyl-CoA within the acetoacetyl-CoA module. a) Schematic of the alternative candidates for this module, which was tested *in vivo* using fructose as substrate in the *ΔphaCAB* strain. The strain was additionally expressing *Cn-fadB’* and *hEc-ydiI* under the control of an L-arabinose inducible promoter. The endogenous route (Cn-PhaA) is driven by the condensation of two acetyl-CoA (Ac-CoA) molecules into acetoacetyl-CoA (AcAc-CoA). This step is thermodynamically challenging with a ∆_r_G^′m^ > 0. An alternative route with a ∆_r_G^′m^ < 0 utilizes the carboxylation of acetyl-CoA into malonyl-CoA upon ATP hydrolysis (Cn-Acc), followed by a decarboxylative Claisen condensation of malonyl-CoA to another acetyl-CoA molecule. This last step requires the heterologous expression of the *nphT7* gene, derived from *Streptomyces* spp. b) Test of the different module variants under heterotrophic conditions, using 1 mM L-arabinose to induce gene expression. Each data point represents the measurement of a biological replicate. The bars indicate the standard deviation calculated on a dataset involving biological duplicates or triplicates.

To test the bypass, we replaced *Cn-phaA* in the L-arabinose inducible plasmid containing *Cn-fadB’* and *hEc-ydiI* with a wildtype and a codon harmonized sequence of *nphT7* from *Streptomyces sp*. (strain CL190), named as *Str-nphT7* and *hStr-nphT7*, respectively. We conjugated the P_BAD__*Str-nphT7_Cn-fadB’_hEc-ydiI and* P_BAD__*hStr-nphT7_Cn-fadB’_hEc-ydiI* plasmids into the Δ*phaCAB* strain. We performed cultivation on fructose as setup for the test. Indeed, upon induction with 1 mM of L-arabinose, the crotonate titers obtained from the strain expressing *hStr-nphT7* improved about three-fold compared to the one expressing *Cn-phaA*, reaching a value of 0.95 ± 0.12 mM (*Fig. 6b*). In contrast, the strain producing the wildtype sequence of *Str-nphT7* resulted in poor crotonate synthesis. In summary, when catalytically functional, the ATP-driven bypass of PhaA is a valuable alternative for increasing flux towards crotonate.

### 3.3 Demonstrating formatotrophic production of crotonate via fed-batch fermentation

After engineering a functional crotonate production pathway using fructose as a substrate, we studied the ability of *C. necator* to synthesize crotonate using formate as feedstock. For this assessment, we used the abovementioned fed-batch system (pH stat) developed in the mini bioreactor setup. We tested both Δ*phaCAB* strains previously characterized for the endogenous conversion of acetyl-CoA to acetoacetyl-CoA (Cn-PhaA) and the new malonyl-CoA bypass (hStr-NphT7, **Fig. 7**). For the sake of simplicity, we will refer to these two strains as the “direct” and the “bypass” strains, respectively (**Fig. 7a**).

**Figure 7.**
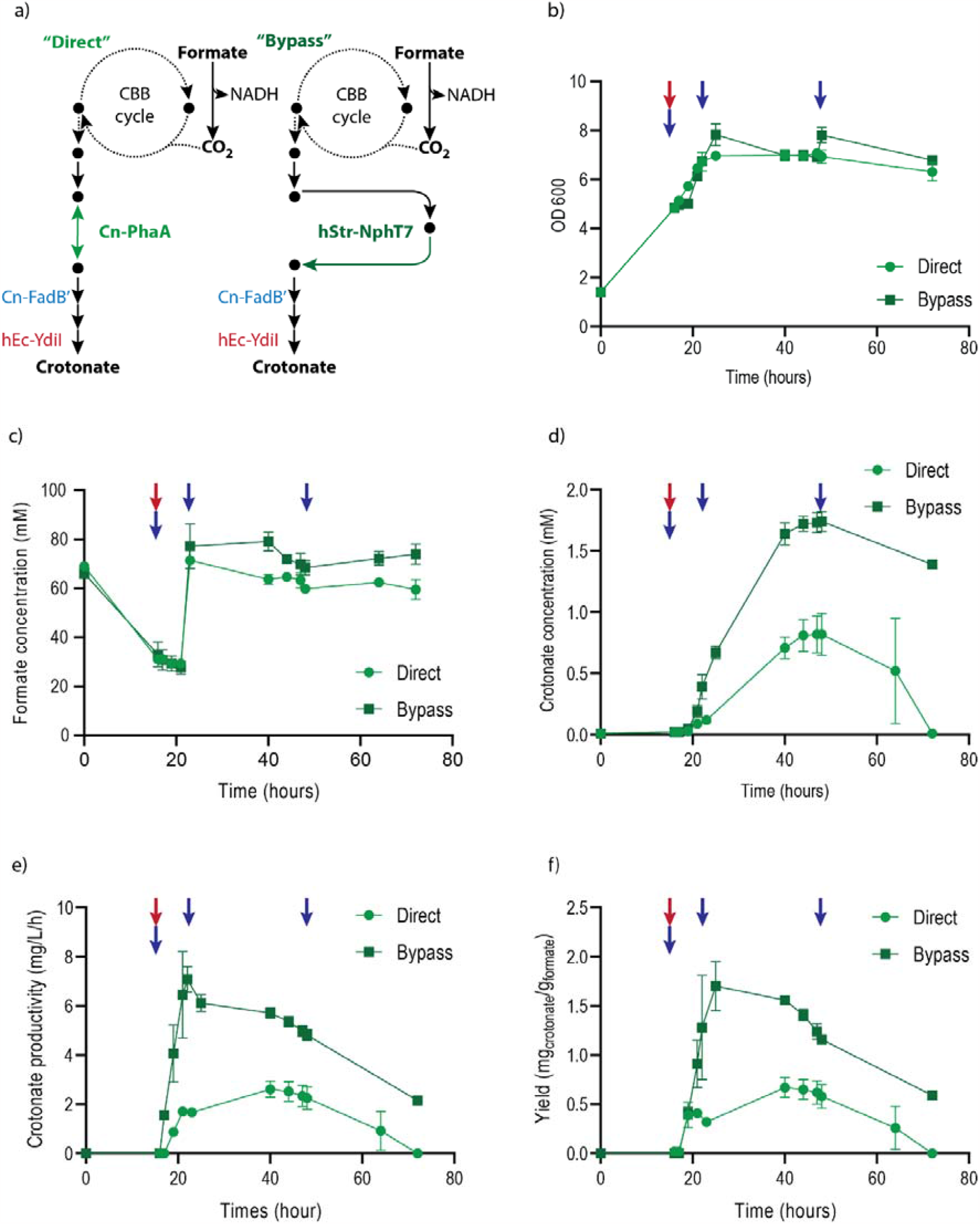
Formatotrophic production of crotonate by the *ΔphaCAB* strains with the linear conversion of acetyl-CoA to acetoacetyl-CoA (PhaA) and the malonyl-CoA bypass (NphT7) variant. a) Schematics representing the lumped formatotrophic metabolism of the “direct” and “bypass” strains, harboring a plasmid-born crotonate biosynthetic pathway expressing *Cn-phaA* or *hStr-nphT7*, respectively. b) Biomass concentration expressed as OD_600_; c) formate concentration in culture media; d) crotonate titers in the medium; e) crotonate volumetric productivity and f) yield of crotonate on formate, calculated from the induction. The red arrows indicate the addition of 1 mM of arabinose (induction of the expression of the crotonate biosynthetic pathway) and the blue arrows the addition of ammonium sulfate (NH_4_)_2_SO_4_: 10 mM at induction; 5 mM at 24 h and 5 mM at 44 h. At 21 h, the concentrations of formate were set to 80 mM by addition of 50 mM of sodium formate. Experiments were performed in duplicates.

We assessed the formatotrophic production of crotonate in fed-batch cultivations. Culture performances are summarized in **Table 3.** During the first 17 hours, both strains grew similarly using formate as only carbon and energy sources (**Fig. 7b**). Both strains grew from OD_600_ 1.5 to about OD_600_ 5.0 (µ=0.08 h^-1^). Despite the addition of formic acid to control the pH (pH stat setup), the concentrations of formate dropped from 70 mM to 30 mM in both cultures before being adjusted back to 80 mM (**Fig. 7c**). This could have been caused by the assimilation of ammonium ions for biomass production resulting in medium acidification.

**Table 3.** Summary of the formatotrophic culture performances of the “direct” *ΔphaCAB* (PhaA) and “linear” *ΔphaCAB* (NphT7) strains.

| Pathway | Unit | Direct strain<br>$\Delta phaCAB$<br>$P_{BAD\_Cn-phaA\_Cn-fadB'_hEc-ydII}$ | Bypass strain<br>$\Delta phaCAB$<br>$P_{BAD\_hStr-nphT7\_Cn-fadB'_hEc-ydII}$ |
| --- | --- | --- | --- |
| Total fermentation time | [h] | 72 | 72 |
| Final OD<sub>600</sub> |  | 7.1 (0.2) | 7.8 (0.5) |
| CDW* | [g<sub>Biomass</sub>/L] | 2.6 (-0.1) | 2.8 (-0.2) |
| Formate consumed | [M] | 5.7 (0.1) | 5.8 (0.1) |
| Crotonate titer max | [mM] | 0.82 (0.15) | 1.74 (0.08) |
|  | [mg<sub>Crotonate</sub>/L] | 69.9 (12.7) | 148.0 (6.8) |
| Biomass yield max | [g<sub>CDW</sub>/mol<sub>FA</sub>]** | 1.50 (0.09) | 1.31 (0.11) |
|  | [mol<sub>CDW</sub>/mol<sub>Formate</sub>] | 0.06 (0.00) | 0.05 (0.00) |
| Crotonate yield max | [mmol<sub>Crotonate</sub>/mol<sub>Formate</sub>] | 0.46 (0.06) | 2.43 (1.11) |
|  | [mg<sub>Crotonate</sub>/g<sub>Formate</sub>] | 0.67 (0.10) | 1.70 (0.25) |
| Crotonate productivity | [mg/(L·h)] | 2.61 (0.33) | 7.08 (0.52) |
| | [μmol/(L·h)] | 31 (4) | 83 (6) |

\* Using a ratio CDW/OD<sub>600</sub> = 0.363 g/L (Grunwald et al., 2015)
\*\* Using biomass composition: CH<sub>1.77</sub>O<sub>0.49</sub>N<sub>0.25</sub> with 3.92% ashes (Aragao et al., 1996; Grousseau, 2012) resulting in a molecular weight =25.35 g/Cmole biomass.
In brackets are the standard deviations

We induced the expression of the crotonate biosynthetic pathway (P_BAD_ promoter) using 1 mM L-arabinose after 17 h. As mentioned above, L-arabinose is not consumed by *C. necator* (Alagesan et al., 2018; Pan et al., 2021), and therefore was not expected to be converted to crotonate. To prevent excessive acidification of the medium upon ammonia assimilation for biomass build-up, we spiked the culture with ammonium sulfate throughout the cultivation (5-10 mM) at 18, 24, and 44 h. The first traces of crotonate were observed 2 hours after the induction for both strains (10 µM for the strain with the linear conversion of acetyl-CoA to acetoacetyl-CoA, and 30 µM for the malonyl-CoA bypass variant (**Fig. 7d**). Between the inductions (17 h) and 24 h, we observed concomitant growth and crotonate production. The linear strain grew up to OD_600_ 7.1 (0.05 h^-1^) and the bypass strain up to OD_600_ 8.0 (µ=0.06 h^-1^). After 24 h, the OD remained relatively stable in all cultures whereas the crotonate titers continued to rise. Maximal crotonate titers were reached after 45 hours in all cultures (**Fig. 7d**).

We therefore calculated the important benchmarks for biotechnological processes, also known as TRY values: titers (in mM); rates -or volumetric productivities- (in mg/(L·h) or µmol/(L·h)); and yields (in mmol_Crotonate_/mol_Formate_). The maximum crotonate titer reached by the direct strain was 0.82 ± 0.15 mM (or 69.9 ± 12.7 mg/L) while the bypass strain reached up to 1.74 ± 0.08 mM or 148.0 ± 6.8 mg/L (**Fig. 7d**). These data confirm the superiority of the pathway with the conversion of acetyl-CoA to acetoacetyl-CoA via the malonyl-CoA bypass. These observations are reflected also in the analysis of the other parameters. We observed a maximal crotonate volumetric productivity during the first 5 to 6 hours after the induction (**Fig. 7e**). In this time interval, the direct strain produced about 2.61 ± 0.33 mg/(L·h) or 31 ± 4 µmol/(L·h) crotonate, whereas the bypass strain produced 7.08 ± 0.52 mg/(L·h) or 83 ± 6 µmol/(L·h) crotonate (**Fig. 7e**). Regarding the crotonate yields (**Fig. 7e**), the bypass strain showed a yield on formate of 2.43 ± 1.11 mmol_Crotonate_/mol_Formate_ 8 hours after the induction (**Fig. 7e**). In the meantime, the maximal crotonate yield on formate of the direct strain was only of 0.46 ± 0.06 mmol_Crotonate_/mol_Formate_ (**Fig. 7e**).

In all cultures, we observed crotonate reassimilation starting after 45 hours of cultivation (**Fig. 7d**). This might result from the induction of crotonate assimilating pathways by *C. necator*. Having a lower crotonate productivity, the direct strain had all its crotonate consumed after 72 hours of cultivation. Instead, in the same time window the bypass strain only partially consumed the produced crotonate, thereby highlighting its superiority as production strain. Nevertheless, all the crotonate was consumed after 100 hours also for this strain (data not shown).

## 4. Discussion

In this work, we combined the optimization of a formatotrophic cultivation setup to the metabolic engineering of *C. necator* to demonstrate formate conversion into the value-added compound crotonate. This is one of the few works describing the production of a platform chemical using formate as only carbon and energy source (**Table 1**). These results add to previous reports on microbial biobased synthesis of crotonate (Dellomonaco et al., 2011; Kim et al., 2016; Liu et al., 2015; Schada Von Borzyskowski et al., 2018; Wang et al., 2019), opening an unprecedented setup for its production directly from formate.

Assessment of formate toxicity revealed an optimum range of concentration between 40 to 90 mM to support high growth-rates and -yields. However, when cultivated in a batch setup in this concentration range, *C. necator* did not reach biomass concentrations above OD_600_ 1.0. A pH stat fed-batch setup -a strategy also recently described elsewhere (Calvey et al., 2023)- revealed to be a successful setup to obtain higher biomass concentrations (g/L range), while supplementing formate within the optimum range. In fact, titrating the medium pH using formic acid allowed to tackle two obstacles at once: first, addition of the acid counterbalanced the alkalinization of the broth; second, it provided additional formate in the bioreactor, which could be assimilated by *C. necator*.

Although our cultivation strategy allowed us to successfully perform strain characterization, few bioprocess-related issues remained. We observed that the formate concentration in the culture broth decreased despite the continuous addition of formic acid requiring regular pulse of sodium formate to maintain cell growth. We speculate that this is caused by ammonia (NH_4_^+^) consumption and of the CO_2_ generation which acidifies the culture medium. In the 2-L bioreactor cultivation described at the beginning of the manuscript, these effects could be mitigated by automatically feeding ammonia in parallel to formic acid (ratio 1 molecule of ammonia per 40 molecules of formic acid) and by maintaining the CO_2_ partial pressure around 5% during the culture. In the 250-mL mini bioreactor setup we could not implement such an automated strategy due to the simpler cultivation setup: this can be a drawback if aiming at obtaining high cell densities in this semi-controlled setup.

Adopting a strategy of modular pathway engineering allowed us to sequentially compare different module variants, thereby assembling the best enzyme arrangement supporting crotonate production. We combined the use of cell lysates and *in vivo* screening (using fructose as substrate) for assessing the different modules. The top performing pathway structure was constructed under the control of an inducible P_BAD_ promoter (L-arabinose) and included a malonyl-CoA bypass for the generation of acetoacetyl-CoA from acetyl-CoA. This architecture fits well with the requirement of a formatotrophic growth mode, where oxidation of formate via Fdh increases the NADH pool, with consequent lower demand of flux through the TCA cycle *via* acetyl-CoA. Therefore, the use of this bypass (at the cost of one ATP investment) confirmed to be a valuable strategy for circumventing thermodynamic bottlenecks when the intracellular pool of acetyl-CoA is limited (Lan and Liao, 2012; Orsi et al., 2022). Under formatotrophic growth conditions, the malonyl-CoA bypass produced almost two-fold higher titers, three-fold higher rates, and five-fold higher yields than the direct route from acetyl-CoA to acetoacetyl-CoA. This evidence demonstrates that tailoring metabolic engineering strategies to formatotrophic growth and not simply adapting the ones already existing for heterotrophic cultivations can significantly improve the bioproduction capacity of the host.

Despite the encouraging results of this proof-of-concept, the current performance (TRY) parameters are still too low for industrial production (**Table 3)**. Further pathway optimization and systems-level metabolic engineering, as well as further bioprocess optimization can probably further boost these performance parameters (Aslan et al., 2017; Lee and Kim, 2015; Meadows et al., 2016; Yim et al., 2011). To better investigate the carbon flux from substrate to product, we performed all cultivations exclusively on a minimal media. This allowed us to gather useful knowledge on the pathway limitations and aided us in identifying a roadmap for further improvements. Building on that, we suggest some metabolic engineering strategies for further improving the crotonate production and thereby getting closer to a consolidated bioprocess.

Starting from crotonate reuptake by *C. necator*, the deletion of competitive consumption pathways could result in a significant improvement of the final product titer. In a similar fashion, deletion of the operon *acoABC* harboring genes encoding for enzymes responsible for acetoin uptake enabled to reach titers of 44 mM of acetoin under autotrophic regimes (Windhorst and Gescher, 2019). Since uptake and consumption of crotonate is likely to overlap with β-oxidation, a route that contains many redundant genes in *C. necator* (Riedel et al., 2014), we recommend analyzing the proteome or the transcriptome of this bacterium when growing on crotonate. This would help in pinpointing those candidates as potential knock-out targets to limit crotonate uptake by the production strain.

Previous work characterizing YdiI for crotonate synthesis in *E. coli* reported a *K*_M_ for crotonyl-CoA higher than 500 µM (Kim et al., 2016). This work showed that despite increased flux towards crotonyl-CoA, the highest crotonate yield on fructose (*Y*_P/S_) measured in test tubes was of only 0.02 mol_Crotonate_/mol_Fructose_. This suggests that YdiI might be catalytically limited also in *C. necator*. This hypothesis is corroborated by the high amounts of 3-HB secreted in the spent medium (**Fig. 4d)**. Therefore, protein engineering of YdiI for improving its specificity towards crotonyl-CoA might be required as next optimization step towards improving crotonate biosynthesis.

One of the possible crotonate biosynthetic routes that we explored involved a partial overlap with the PHB pathway. In fact, this route is known to be highly functional in *C. necator*, where it can reach up to 80% w/w of the whole CDW (Riedel et al., 2014). Nevertheless, overexpression of *Cn*-*phaB* did not result in significant improvements of crotonate synthesis (*Fig. 4c*). This is probably because no endogenous *C. necator* enzyme could catalyze the dehydration step from 3-(*R*)-HB-CoA to crotonyl-CoA. To fully harness the NADPH-dependent route for crotonate biosynthesis, one would have to express dehydratase genes specific for 3-(*R*)-HB-CoA such as *e*.*g*., *phaJ* from *Rhodospirillum rubrum* (Reiser et al., 2000), or *croR* from *M. extorquens* (Schada Von Borzyskowski et al., 2018).

Another potential bottleneck to tackle in crotonate biosynthesis might involve the increase in precursors supply. Generation of acetyl-CoA from the central metabolism involves decarboxylation of pyruvate, with consequent loss of CO_2_ for product formation. Therefore, additional system-level strategies could be explored. As recently proposed, employing a heterologous phosphoketolase could be a useful option for further improving the acetyl-CoA pool when growing on formate, where a xylulose 5-phosphate phosphoketolase (Xpk) activity could be used to source additional acetyl-CoA directly from the CBB cycle (Janasch et al., 2022).

Improving formate assimilation in *C. necator* could also positively impact production capacity. Recently, the improvement of the formatotrophic growth-rate of *C. necator* through the endogenous CBB cycle was reported (Calvey et al., 2023). Using such a strain could improve the volumetric productivity (g/L/h) of crotonate, as well as reducing the duration of the reactor run (operating costs). Alternatively, synthetic formate assimilation routes can be explored. The reductive glycine pathway, known as the most ATP-efficient formate assimilation route (Bar-Even et al., 2013), has already been implemented (Claassens et al., 2020) and was then further optimized to exceed the biomass yield of the Calvin cycle (Dronsella et al., 2022) in this bacterium. Therefore, we anticipate that using a more efficient formate assimilation route could in the future positively impact product yields.

We believe that this result will encourage further investigation on the biological conversion of formate into value-added products for the bioeconomy. Moreover, we pose that by following a holistic approach towards the optimization of *C. necator*’s metabolic network it will be possible to tailor its performance to the requirements of the formatotrophic cultivation setup we developed in this work. Once these challenges are tackled, it will be possible to significantly advance this proof-of-concept into a highly sustainable bioprocess based on truly renewable substrates.

## Acknowledgments

We dedicate this work to the memory of Arren Bar-Even, who was involved in the initial conceptualization of this paper, as well as the funding acquisition part. We thank Pablo I. Nikel and Sebastian Wenk for their feedbacks and critical reading of this manuscript, and Stefano Donati for helpful discussion and suggestions. We also thank Nicolò Baldi and Seohyoung Kim for helping to design the thioesterase activity assay. F.C., B.D., F.K. and E.O. were financially supported by the German Ministry of Education and Research (BMBF) through the grant TRANSFORMATE (033RC023) within the CO2-WIN program. Moreover, E.O. acknowledges financial support from The Novo Nordisk Foundation through grants NNF20CC0035580 and NNF21OC0070572. N.J.C. acknowledges support from a Veni grant (VI.Veni.192.156) from the Dutch Science Organization (NWO).

## CRediT authorship contribution statement

*Florent Collas*: Conceptualization, Methodology, Formal Analysis, Investigation, Writing – Original Draft, Visualization; *Beau Dronsella*: Conceptualization, Formal Analysis, Investigation, Writing – Original Draft, Writing – Review & Editing; *Armin Kubis*: Conceptualization, Writing – Original Draft; *Karin Schann*: Formal Analysis, Investigation, Visualization; *Sebastian Binder*: Formal Analysis, Investigation; *Nils Arto*: Formal Analysis, Investigation; *Nico J. Claassens*: Conceptualization, Supervision, Writing – Original Draft, Writing – Review & Editing; *Frank Kensy*: Conceptualization, Supervision, Project Administration, Funding Acquisition, Writing – Original Draft; *Enrico Orsi*: Conceptualization, Methodology, Formal Analysis, Investigation, Writing – Original Draft, Visualization, Writing – Review & Editing.

