## Supplementary for "Engineering the biological conversion of formate into crotonate in *Cupriavidus necator*"

### **Supplementary material**

### **Tables**

#### **Table S1. Primers used in this study**

| **Primer** | **Sequence (5´-3´)** | **Description** |
| --- | --- | --- |
| 5483 | CCAAGATGGATCTGGTGAGG | Forward internal primer to cas9 on the pCas9 plasmid. Used to confirm plasmid loss in ∆*phaCAB C. necator* strains |
| 5486 | GTTAACACCGAGATTACCAAAG | Reverse internal primer to cas9 on the pCas9 plasmid. Used to confirm plasmid loss in ∆*phaCAB C. necator* strains |
| 5549 | TTCGAGCTCGGTACCCGGGGATCCTCTAGAAAAAACCGTTATTGACACAGGTGGAAATTTAGAATATACTGTTAGTAA | To use with 5550 to create Pj5 promoter |
| 5550 | ATGTATATCTCCTTCGGAGCTCTTACTAACAGTATATTCTAAATTTCCACCTG | To use with 5549 to create Pj5 promoter. Includes RBS1 |
| 5624 | TCTAGAGGATCCCCGGGTAC | Forward primer to linearize pSEVA221 and open its MCS |
| 5625 | GTCGACCTGCAGGCATG | Reverse primer to linearize pSEVA221 and open its MCS |
| 6679 | AGAATATACTGTTAGTAAgagctcCGAAGGAGATATACATatgcatcatcaccatcaccacatatg | Forward primer to amplify *Ec-ydiI*, including RBS1 and overhang for Pj5 |
| 5556 | GGCCGCAAGCTTGCATGCCTGCAGGTCGACtcacaaaatggcggtcgtcaatc | Reverse primer to amplify *Ec-ydiI*. Contains overhang for MCS of pSEVA221 |
| 6093 | gtatctggctgctgacCGG | Forward primer to amplify the synthetic construct containing RBS4 and hEc-ydiI |
| 6094 | GGCCGCAAGCTTGCATG | Reverse primer to amplify the synthetic construct containing *hEc-ydiI*. It anneals on MCS of pSEVA221 |
| 6680 | GTTAGTAAgagctcCGAAGGAGATATACATATGATTTGGAAGCGGAAGATTACGC | Forward primer to amplify h*Ec-ydiI*, including RBS1 and overhang for Pj5 |
| 6681 | GTTAGTAAgagctcCGAAGGAGATATACATATGAGTCAGGCGCTAAAAAATTTACTGAC | Forward primer to amplify *Ec-tesB*, including RBS1 and overhang for Pj5 |
| 6682 | GGCCGCAAGCTTGCATGCCTGCAGGTCGACTTAATTGTGATTACGCATCACCCCTTC | Reverse primer to amplify *Ec-tesB*. Contains overhang for MCS of pSEVA221 |
| 5551 | AGAATATACTGTTAGTAAgagctcCGAAGGAGATATACATATGACTGACGTTGTCATCGTATCC | Forward primer to amplify *Cn-phaA*, including RBS1 and overhang for Pj5 |
| 5552 | CCTTGTTCTCCTTTGgaattctacgattacTTATTTGCGCTCGACTGCCAGC | Reverse primer to amplify *Cn-phaA*, including overhangs for RBS5. |
| 5679 | gtggcgctggcagtcgagcgcaaataagtaatcgtagaattcCAAAGGAGAACAAGGAtgcatcatcaccatcaccacctttacaaaggc | Forward primer for *Ec-fadB* amplification. Ist overhang anneals with the 3' end of phaA, and includes the sequence for RBS5. |
| 5554 | ACTTACTCCTCCGgtcagcagccagatacttaagccgttttcaggtcgccaa | Reverse primer for the amplification of *Ec-fadB*. Includes the overhangs for RBS4. |
| 5770 | ggcgacctgaaaacggcttaagtatctggctgctgacCGGAGGAGTAAGTatgcatcatcaccatcaccacatatggaaacgg | Forward primer for cloning *Ec-ydiI* in the last position of the operon. Includes overhangs for RBS4. |
| 5771 | gtgctgttcgtcacgattgacgaccgccattttgtgaGTCGACCTGCAGGCATGcaagcttgcg | Reverse primer for cloning *Ec-ydiI* in the last position of the operon. Includes overhangs for the MCS of pSEVA221 |
| 6088 | CCTTGTTCTCCTTTGgaattctacgattac | Forward primer to linearize the crotonate plasmid, with removal of the gene in position 2 in the operon |
| 6089 | GTATCTGGCTGCTGACCGG | Reverse primer to linearize the crotonate plasmid, with removal of the gene in position 2 in the operon |
| 6091 | gtaatcgtagaattcCAAAGGAGAACAAGG | Forward primer to amplify the synthetic construct *hEc-fadB*. Include overhangs for annealing with the linearized plasmid, and introduce the gene in the position 2 of the operon, under the control of RBS5 |
| 6092 | TACTTACTCCTCCGgtcagc | Reverse primer to amplify the synthetic construct *hEc-fadB*. Include overhangs for annealing with the linearized plasmid, and introduce the gene in the position 2 of the operon |
| 6248 | gtaatcgtagaattcCAAAGGATGTCCAATTTCATCGTCAAGAAGGTC | Forward primer to amplify the synthetic construct *Cn-fadB'*. Include overhangs for annealing with the linearized plasmid, and introduce the gene in the position 2 of the operon, under the control of RBS5 |
| 6096 | ACTCCTCCGgtcagcagccagatacTTAGTTACGCACCGGCTTGC | Reverse primer to amplify the synthetic construct *Cn-fadB'*. Include overhangs for annealing with the linearized plasmid, and introduce the gene in the position 2 of the operon |
| 6249 | gtaatcgtagaattcCAAAGGatgacagcgcagtatcaggttc | Forward primer to amplify the synthetic construct *Cn-fadB1*. Include overhangs for annealing with the linearized plasmid, and introduce the gene in the position 2 of the operon, under the control of RBS5 |
| 6098 | ACTCCTCCGgtcagcagccagatacttaaccgttgaagcccttgcc | Reverse primer to amplify the synthetic construct *Cn-fadB1*. Include overhangs for annealing with the linearized plasmid, and introduce the gene in the position 2 of the operon |
| 7578 | AgtaatcgtagaattcCAAAGGAGAACAAGGATGACTCAGCGCATTGCGTATG | Forward primer to amplify the synthetic construct *Cn-phaB1*. Include overhangs for annealing with the linearized plasmid, and introduce the gene in the position 2 of the operon, under the control of RBS5 |
| 7579 | ACTTACTCCTCCGgtcagcagccagatacTCAGCCCATATGCAGGCCG | Reverse primer to amplify the synthetic construct *Cn-phaB1*. Include overhangs for annealing with the linearized plasmid, and introduce the gene in the position 2 of the operon |
| 7437 | CTCGGTACCCGGGGATCCTCTAGAATTATGACAACTTGACGGCTACATC | Forward primer to amplify the PBAD promoter, including araC, from E. coli genomic DNA |
| 7438 | CATATGTATATCTCCTTCGgagctcTcaaaattatttctagagggaaaccgttgtggtctccctatggagaaacagtagagagttgc | Reverse primer to amplify the PBAD promoter, including araC, from E. coli genomic DANN. Contains overhangs for RBS1 on pSEVA plasmid |
| 7439 | AGAGCTCCGAAGGAGATATACATATG | When combined with 5624, it serves as primer for linearizing the crotonate plasmid, and exclude the promoter Pj5. It also anneals |
| 6281 | ATGTATATCTCCTTCGgagctc | Forward primer to use for linearizing the crotonate plasmid, excluding gene in position 1. |
| 6282 | gtaatcgtagaattcCAAAGGAG | Reverse primer to use for linearizing the crotonate plasmid, excluding gene in position 1. |
| 7611 | GTAAgagctcCGAAGGAGATATACATATGCATCATCACCATCACCACAC | Forward primer to amplify *Str-nphT7*, and clone it in position 1 of the operon, under the control of PBAD promoter and RBS1 |
| 7612 | CTCCTTTGgaattctacgattacGCTAGCTCTAGATTACCATTCGATC | Reverse primer to amplify *Str-nphT7*, and clone it in position 1 of the operon. Contains overhang for RBS5. |
| 6956 | GTAAgagctcCGAAGGAGATATACATATGACtGAtGTCCGATTCCGC | Forward primer to amplify *hStr-nphT7*, and clone it in position 1 of the operon. |
| 6957 | TTCTCCTTTGgaattctacgattacCTACCACTCGATCAGaGCG | Reverse primer to amplify *hStr-nphT7*, and clone it in position 1 of the operon. |
| 6217 | GAGAATCGCCGTCGTGAG | To sequence *hEc-ydiI* |
| 3117 | CGGTCTCTCCGGGCTATATC | To sequence *Cn-phaB* |
| 3425 | CCTTGGCGGTGGAGTAGTTG | To sequence *Cn-phaB* |
| 3058 | AGTGGCTGGAGCAGCAGAAG | To sequence *Cn-phaB* |
| 7128 | GGCaGCTCCCGTGTTGC | To sequence *hStr-nphT7* |
| 7002 | CGAGGTaGCCTGGTCGTC | To sequence *hStr-nphT7* |
| 7181 | GGTACGGGTGCCTACG | To sequence *hStr-nphT7* |
| 7182 | GGCaGCTCCCGTGTTGC | To sequence *hStr-nphT7* |
| 5648 | CGCTGGACAGCATGTCC | To sequence *Cn-phaA* |
| 6100 | CAAGGAACTGGGCCTGACC | To sequence *Cn-phaA* |
| 6338 | AGTGGACCCCGCAAGACCTG | To sequence *Cn-phaA* |
| 5511 | GGCAAAGGCCTCGTTGAT | To sequence *Cn-phaA* |
| 6101 | GTTAAGTCAGTGGCTGCACTTTG | To sequence *Ec-fadB* |
| 6102 | CGATGGTCTGAAACTGGCTGG | To sequence *Ec-fadB* |
| 6103 | gcgttccgctggctg | To sequence *Ec-fadB* |
| 6104 | GCGCATGGACTGGAAGCAC | To sequence *Cn-fadB'* |
| 6105 | GCGATCCGCTGGGGC | To sequence *Cn-fadB'* |
| 6106 | GAAGCGCGCCAGATGG | To sequence *Cn-fadB'* |
| 6107 | TCGGTCGCCATGGGTG | To sequence *Cn-fadB1* |
| 6108 | tcatcgaggccgtgttcgaag | To sequence *Cn-fadB1* |
| 6109 | gatgaagcgctacgccaagg | To sequence *Cn-fadB1* |

#### **Table S2. Codon harmonized gene sequences used in this study**

| **Gene name** | **Harmonized sequence (5'-3')** |
| --- | --- |
| *hEc-ydiI* | ATGATTTGGAAGCGGAAGATTACGCTCGAGGCGCTCAACGCAATGGGGGAGGGGAATATGGTCGGGTTTCTCGACATTCGGTTCGAGCACATTGGGGACGATACGCTCGAGGCGACGATGCCCGTCGATTCCCGGACGAAACAGCCCTTTGGGCTCCTCCACGGGGGGGCGTCCGTCGTCCTCGCGGAGTCCATTGGGTCCGTCGCGGGGTACCTCTGCACGGAGGGGGAACAAAAGGTCGTCGGGCTCGAGATTAACGCAAATCATGTCCGGTCCGCGCGTGAGGGGCGGGTCCGGGGGGTCTGTAAGCCCCTCCACTTGGGGTCCCGGCATCAGGTCTGGCAGATTGAGATTTTTGACGAAAAGGGGCGGCTCTGTTGCTCCTCCCGTCTCACGACGGCGATTCTCTAA |
| *hEc-fadB* | ATGACtGAtGTCCGATTCCGCATTATCGGTACGGGTGCCTACGTACCGGAACGGATCGTCTCCAACGATGAAGTCGGtGCaCCaGCaGGtGTtGACGACGACTGGATCACCCGCAAGACCGGTATCaGaCAGCGTCGCTGGGCaGCtGACGACCAGGCtACCTCGGACCTGGCCACGGCtGCaGGGCGGGCAGCaCTGAAAGCGGCGGGCATCACGCCCGAGCAGCTGACCGTGATCGCGGTCGCCACCTCCACGCCGGACCGGCCGCAGCCGCCCACGGCGGCCTATGTCCAGCACCACCTCGGTGCGACCGGCACTGCtGCaTTCGACGTCAACGCtGTCTGCTCCGGtACCGTGTTCGCGCTGTCCTCGGTGGCtGGtACCCTCGTGTACCGGGGtGGTTACGCtCTGGTCATCGGaGCtGACCTGTACTCGCGCATCCTCAACCCtGCCGACCGtAAGACGGTCGTGCTGTTCGGaGACGGaGCaGGtGCAATGGTCCTCGGaCCGACCTCGACaGGCACGGGaCCtATCGTCCGaCGaGTCGCCCTGCACACCTTCGGtGGaCTCACCGACCTGATCCGTGTGCCCGCGGGaGGtAGCCGCCAGCCaCTGGACACGGATGGaCTCGACGCaGGACTGCAGTACTTCGCGATGGACGGaCGTGAGGTGCGaCGCTTCGTCACGGAGCACCTGCCaCAGCTGATCAAGGGCTTCCTGCACGAGGCCGGtGTCGACGCaGCtGACATCAGCCACTTCGTGCCGCATCAGGCCAACGGTGTCATGCTCGACGAGGTCTTCGGtGAGCTGCATCTGCCGCGGGCGACCATGCACCGGACGGTCGAGACCTACGGCAACACGGGAGCtGCCTCCATCCCGATCACCATGGACGCtGCtGTGCGCGCtGGTTCCTTCCGGCCGGGaGAGCTGGTCCTGCTGGCaGGGTTCGGaGGtGGtATGGCtGCaAGCTTCGCtCTGATCGAGTGGTAG |
| *hStr-nphT7* | ATGACtGAtGTCCGATTCCGCATTATCGGTACGGGTGCCTACGTACCGGAACGGATCGTCTCCAACGATGAAGTCGGtGCaCCaGCaGGtGTtGACGACGACTGGATCACCCGCAAGACCGGTATCaGaCAGCGTCGCTGGGCaGCtGACGACCAGGCtACCTCGGACCTGGCCACGGCtGCaGGGCGGGCAGCaCTGAAAGCGGCGGGCATCACGCCCGAGCAGCTGACCGTGATCGCGGTCGCCACCTCCACGCCGGACCGGCCGCAGCCGCCCACGGCGGCCTATGTCCAGCACCACCTCGGTGCGACCGGCACTGCtGCaTTCGACGTCAACGCtGTCTGCTCCGGtACCGTGTTCGCGCTGTCCTCGGTGGCtGGtACCCTCGTGTACCGGGGtGGTTACGCtCTGGTCATCGGaGCtGACCTGTACTCGCGCATCCTCAACCCtGCCGACCGtAAGACGGTCGTGCTGTTCGGaGACGGaGCaGGtGCAATGGTCCTCGGaCCGACCTCGACaGGCACGGGaCCtATCGTCCGaCGaGTCGCCCTGCACACCTTCGGtGGaCTCACCGACCTGATCCGTGTGCCCGCGGGaGGtAGCCGCCAGCCaCTGGACACGGATGGaCTCGACGCaGGACTGCAGTACTTCGCGATGGACGGaCGTGAGGTGCGaCGCTTCGTCACGGAGCACCTGCCaCAGCTGATCAAGGGCTTCCTGCACGAGGCCGGtGTCGACGCaGCtGACATCAGCCACTTCGTGCCGCATCAGGCCAACGGTGTCATGCTCGACGAGGTCTTCGGtGAGCTGCATCTGCCGCGGGCGACCATGCACCGGACGGTCGAGACCTACGGCAACACGGGAGCtGCCTCCATCCCGATCACCATGGACGCtGCtGTGCGCGCtGGTTCCTTCCGGCCGGGaGAGCTGGTCCTGCTGGCaGGGTTCGGaGGtGGtATGGCtGCaAGCTTCGCtCTGATCGAGTGGTAG |

### **Figures**

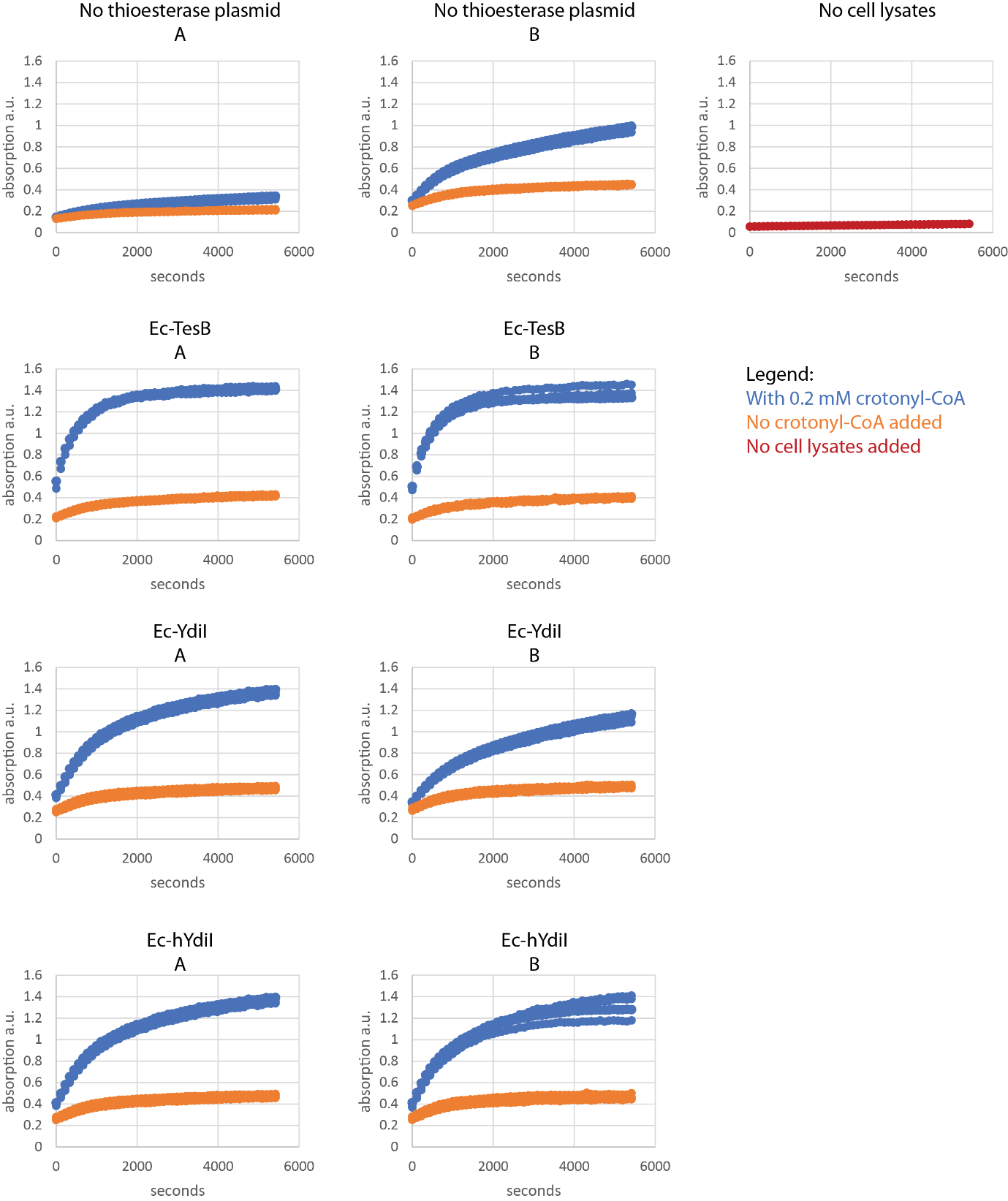

**Figure S1.** Overview of the absorption profiles at 412 nm for the different strains. Two biological replicates for each strain were tested in triplicates. Blue lines correspond to the assay, while orange lines correspond to the negative control (no crotonyl-CoA added). The panel with the red lines shows the change in absorption when crotonyl-CoA is supplemented, but no cell lysates are present.

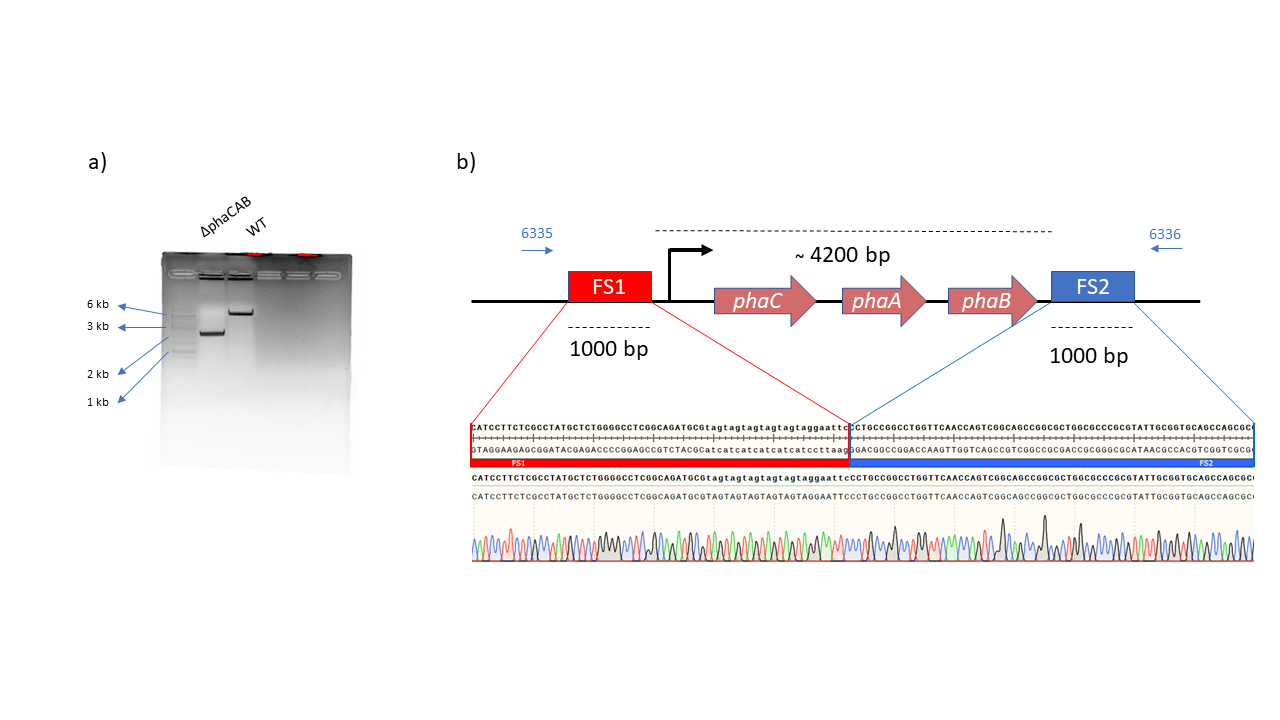

**Figure S2.** Deleted *phaCAB* operon in *Cupriavidus necator*. a) Colony PCR with the primer set 6335/6336, annealing outside the flanking sites FS1 and FS2. b) confirmed deletion of the operon via Sanger sequencing.
